# Shared experience, shared memory: a common structure for brain activity during naturalistic recall

**DOI:** 10.1101/035931

**Authors:** J. Chen, Y.C. Leong, K.A. Norman, U. Hasson

## Abstract

Our daily lives revolve around sharing experiences and memories with others. When different people recount the same events, how similar are their underlying neural representations? In this study, participants viewed a fifty-minute audio-visual movie, then verbally described the events while undergoing functional MRI. These descriptions were completely unguided and highly detailed, lasting for up to forty minutes. As each person spoke, event-specific spatial patterns were reinstated (movie-vs.-recall correlation) in default network, medial temporal, and high-level visual areas; moreover, individual event patterns were highly discriminable and similar between people during recollection (recall-vs.-recall similarity), suggesting the existence of spatially organized memory representations. In posterior medial cortex, medial prefrontal cortex, and angular gyrus, activity patterns during recall were more similar between people than to patterns elicited by the movie, indicating systematic reshaping of percept into memory across individuals. These results reveal striking similarity in how neural activity underlying real-life memories is organized and transformed in the brains of different people as they speak spontaneously about past events.

---

We tend to think of memories as personal belongings, a specific set of episodes unique to each person's own mind. Every person perceives the world in his or her own way and describes the past through the lens of individual history, selecting different details or themes as most important. On the other hand, memories do not seem to be entirely idiosyncratic; for example, after seeing the same list of pictures, there is considerable correlation between people in which items are remembered^1^. The capacity to create and share memories is essential for our ability to interact with others and form social groups. The macro and micro-processes by which shared experiences contribute to a community’s collective memory have been extensively studied across varied disciplines^2–7^, yet relatively little is known about how shared experiences shape memory representations in the brains of people who are engaged in unsupervised natural recollection. If two people freely describe the same event, how similar (across brains) are the neural codes elicited by that event?

Despite our differences, human brains have much in common with one another. Similarities exist not only at the anatomical level, but also in terms of functional organization. Given the same stimulus, an expanding ring for example, regions of the brain that process sensory (visual) stimuli will respond in a highly predictable and similar manner across different individuals. This predictability is not limited to sensory systems: shared activity across people has also been observed in higher-order brain regions (e.g., the default mode network^8^ [DMN]) during the processing of semantically complex real-life stimuli such as movies and stories^9–16^. Interestingly, shared responses in these high-order areas seem to be associated with narrative content and not with the physical form used to convey it^14^. It is unknown – at any level of the cortical hierarchy – to what extent the similarity of human brains during shared perception is recapitulated during shared recollection. This prospect is made especially challenging when recall is spontaneous and spoken, and the selection of details left up to the rememberer (rather than the experimenter), as is often the case in real life.

Although a memory is an imperfect replica of the original experience, the imperfection may serve a purpose. As demonstrated by Jorge Luis Borges in his story “Funes the Memorious”^17^, a memory system that perfectly recorded all aspects of experience, without the ability to compress, abstract, and generalize the to-be-remembered information, would be useless for cognition and behavior. In other words, perceptual representations undergo some manner of beneficial modification in the brain prior to recollection. Therefore, memory researchers can ask two complementary questions: 1) to what extent a memory resembles the original event; and 2) what transformations take place between perceptual experience and later recollection. The first question has been extensively explored in neuroscience; many studies have shown that neural activity during perception of an event is reactivated to some degree during recollection of that event^18–20^. However, the laws governing transformations between percept and recollection are not well understood.

In this paper, we introduce a novel inter-subject pattern correlation framework that reveals shared memory representations and shared memory transformation processes across the brain. Participants watched a movie and then were asked to verbally recount the full series of events, aloud, in their own words, without any external cues. Despite the unconstrained nature of this behavior, we found that spatial patterns of brain activity observed during movie-viewing were reactivated during spoken recall (movie-vs.-recall similarity). The reactivated patterns were observed in an expanse of high-order brain areas that are implicated in memory and conceptual representation, broadly overlapping with the DMN. We also observed that these spatial activity patterns were similar across individuals during spoken recall (recall-vs.-recall similarity) and highly specific to individual events in the narrative (i.e., discriminable), suggesting the existence of a common spatial organization or code for memory representations. Strikingly, in many areas within the DMN, we found that neural representations had transformed between perception and recall in a systematic manner across individuals (recall-vs.-recall similarity was stronger than movie-vs.-recall similarity). This transformation was predictive of subsequent memory for events.

Overall, the results suggest the existence of a common spatial organization or topography for memory representations in the brains of different individuals, concentrated in high-level cortical areas (including the DMN) and robust enough to be observed as people speak freely about the past. Furthermore, neural representations in these brains regions were modified between perceptual experience and memory in a systematic manner across different individuals, suggesting a shared process for beneficial memory transformation.

## RESULTS

**Spontaneous spoken recall**. Seventeen participants were presented with a 50-minute segment of an audio-visual movie (BBC’s “Sherlock”^21^) while undergoing functional MRI (Fig. 1A). They were informed before viewing that they would later be asked to describe the movie. Following the movie, participants were instructed to describe aloud what they recalled of the movie in as much detail as they could, with no visual input or experimenter guidance, during brain imaging. Participants were allowed to speak for as long as they wished, on whatever aspects of the movie they chose, while their speech was recorded with an fMRI-compatible microphone.

**Figure 1.**
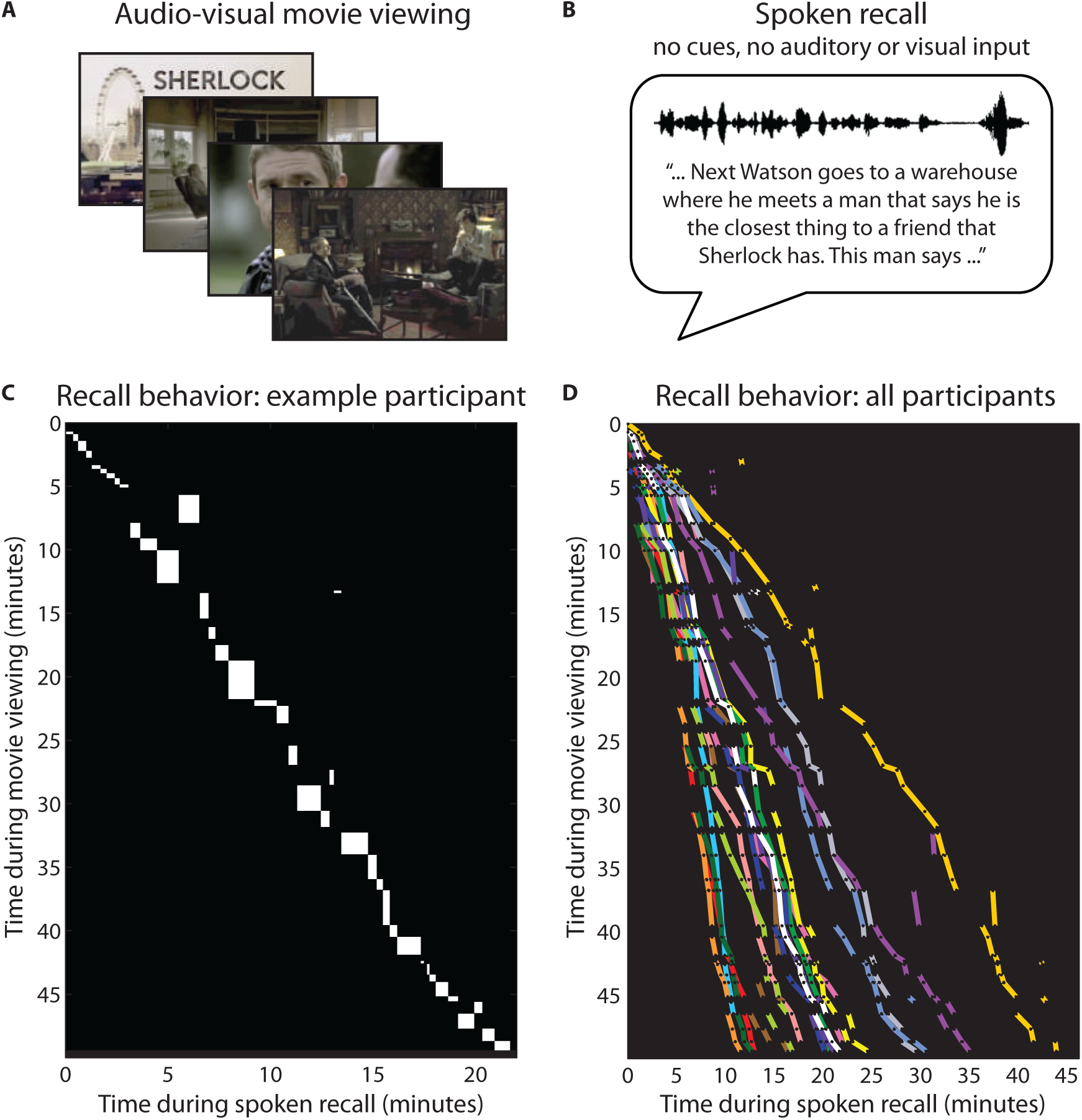
Experiment design and behavior. **A)** In Run 1, participants viewed a 50-minute movie, BBC's Sherlock (episode 1). **B)** In the immediately following Run 2, participants verbally recounted aloud what they recalled from the movie. Instructions to “retell what you remember in as much detail as you can” were provided before the start of the run. No form of memory cues, time cues, or any auditory/visual input were provided during the recall session. Speech was recorded via microphone. **C)** Diagram of scene durations and order for movie viewing and spoken recall in a representative participant. Each rectangle shows, for a given scene, the temporal position (location on y-axis) and duration (height) during movie viewing, and the temporal position (location on x-axis) and duration (width) during recall. **D)** Summary of durations and order for scene viewing and recall in all participants. Each line segment shows, for a given scene, the temporal position and duration during movie viewing and during recall; i.e., a line segment in [D] corresponds to the diagonal of a rectangle in [C]. Each color indicates a different participant (N=17). See also Tables S1, S2.

Without any guidance from the experimenters, participants were able to recall the events of the movie with remarkable accuracy and detail, with the spoken recall sessions lasting on average 21.7 minutes (min: 10.8, max: 43.9, s.d. 8.9) and consisting of on average 2657.2 words (min: 1136, max: 5962, s.d. 1323.6; Table S1). Participants’ recollections primarily concerned the plot: characters’ actions, speech, and motives, and the locations in which these took place. Additionally, many participants described visual features (e.g., colors and viewpoints) and emotional elements (e.g., characters’ feelings). The movie was divided into 50 “scenes” (11 – 180 [s.d. 41.6] seconds long), following major shifts in the narrative (e.g., location, topic, and/or time, as defined by an independent rater; see Experimental Procedures). The same “scenes” were identified in the auditory recordings of the recall sessions based on each participant’s speech. On average, 34.4 (s.d. 6.0) scenes were successfully recalled. A sample participant’s complete recall behavior is depicted in Fig. 1C; see Fig. 1D for a summary of all participants’ recall behavior. Scenes were recalled largely in the correct temporal order, with an average of 5.9 (s.d. 4.2) scenes recalled out of order. The temporal compression during recall (i.e., the duration of recall relative to the movie; see slopes in Fig. 1D) varied widely, as did the specific words used by different participants (see Table S2 for examples).

**Neural reinstatement within participants**. Before examining shared neural patterns across people, we first wished to establish to what extent, and where in the brain, the task elicited similar activity between movie-viewing (encoding) and spoken recall within each participant, i.e., movie-vs.-recall neural pattern reinstatement. Studies of pattern reinstatement are typically performed within-participant, using relatively simple stimuli such as single words, static pictures, or short video clips, often with many training repetitions to ensure successful and vivid recollection of studied items ^20,22–28^. Thus, it was not known whether pattern reinstatement could be measured with fMRI using a single exposure to such an extended complex stimulus and unconstrained spoken recall behavior.

For each participant, brain data were transformed to a common space (MNI) and then the data from movie-viewing and spoken recall were each divided into the same 50 scenes as defined for the behavioral analysis. This allowed us to match time periods during the movie to time periods during recall. All timepoints within each scene were averaged, resulting in one pattern of brain activity, in volume space, for each scene. The pattern for each movie scene (“stimulus-induced pattern”) was compared to the pattern during spoken recall of that scene (“recollection pattern”), within-participant, using Pearson correlation (Fig. 2A). The analysis was performed in a searchlight^29^ across the brain volume (i.e., repeated in 15 × 15 × 15 mm cubes centered on every voxel in the brain). Statistical significance was evaluated using a permutation^30^ analysis that compares the neural pattern similarity between matching scenes against that of non-matching scenes, corrected for multiple comparisons using FDR (q < 0.05, two-tailed, Fig 2A). This analysis reveals regions containing scene-specific reinstatement patterns, as statistical significance is only reached if matching scenes (same scene in movie and recall) can be differentiated from non-matching scenes.

**Figure 2.**
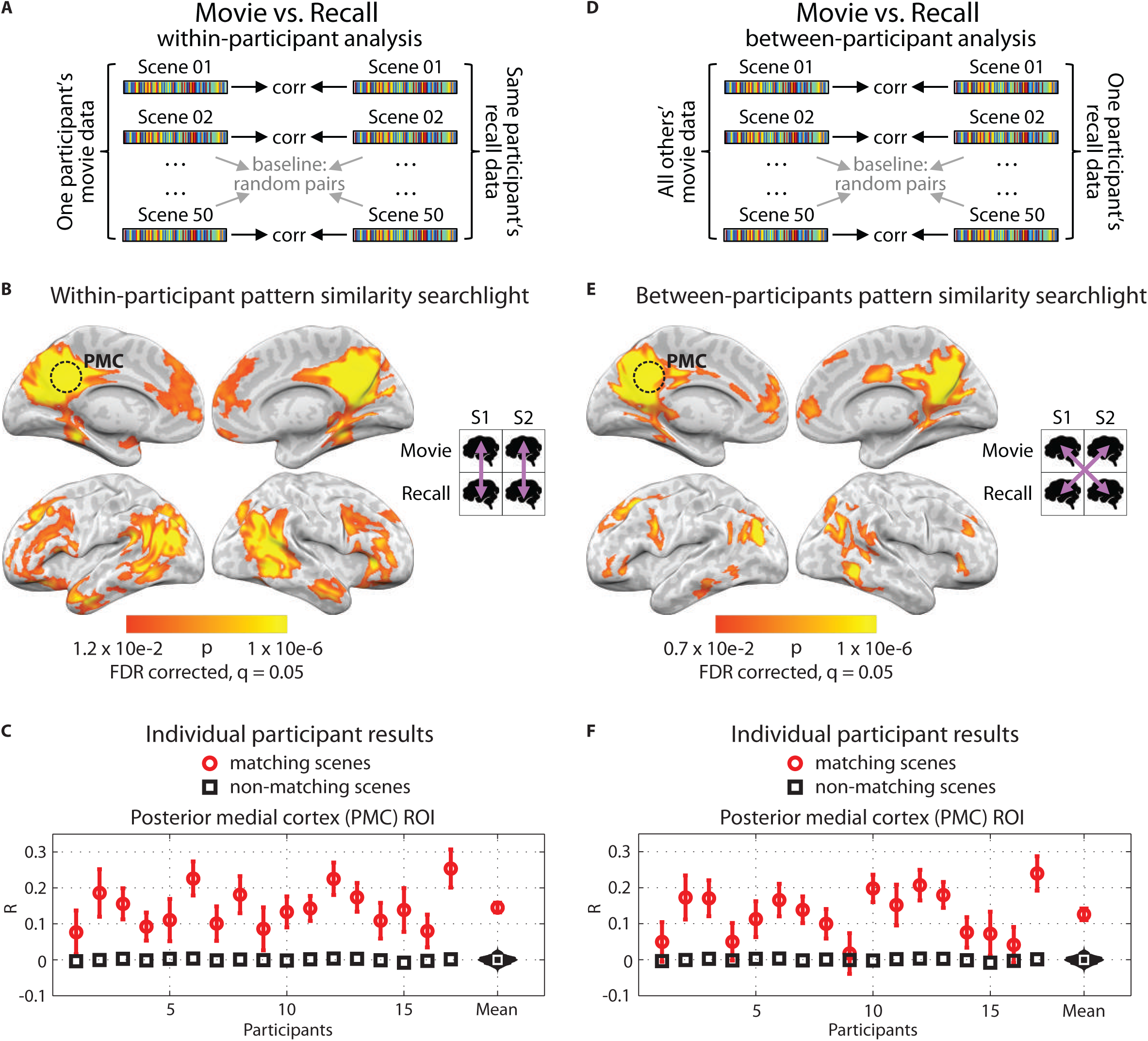
Pattern similarity between movie and recall. **A)** Schematic for within-participant movie-vs.-recall (reinstatement) analysis. BOLD data from the movie and from the recall sessions were divided into scenes, then averaged across time within-scene, resulting in one vector of voxel values for each movie scene and one for each recalled scene. Correlations were computed between matching pairs of movie/recalled scenes within participant. Statistical significance was determined by shuffling scene labels to generate a null distribution of the participant average. **B)** Searchlight map showing where significant reinstatement was observed; FDR correction q = 0.05, p = 0.012. Searchlight was a 5×5×5 voxel cube. **C)** Reinstatement values for all 17 participants in independently-defined PMC (posterior medial cortex). Red circles show average correlation of matching scenes and error bars represent standard error across scenes; black squares show average of the null distribution. At far right, the red circle shows the true participant average and error bars represent standard error across participants; black histogram shows the null distribution of the participant average; white square shows mean of the null distribution. **D)** Schematic for between-participants movie-vs.-recall analysis. Same as [A], except that correlations were computed between every matching pair of movie/recalled scenes between participants. **E)** Searchlight map showing regions where significant between-participants movie-vs.-recall similarity was observed; FDR correction q = 0.05, p = 0.007. **F)** Reinstatement values in PMC for each participant in the between-participants analysis. See also Figures S1, S2.

The searchlight analysis revealed a large set of brain regions in which the scene-specific spatial patterns observed during movie-viewing were reinstated during the spoken recall session (Fig. 2B). These areas included posterior medial cortex, medial prefrontal cortex, parahippocampal cortex, and posterior parietal cortex (Fig. 2B). This set of regions corresponds well with the extensively-studied “default mode network” (DMN)^8,31^, and encompasses areas that are known to respond during cued recollection in more traditional paradigms^32^. Individual participant correlation values for independently-defined posterior medial cortex (PMC) are shown in Fig. 2C. PMC was selected for illustration purposes because the region is implicated as having a long (on the order of minutes) memory-dependent integration window in studies that use real-life stimuli such as movies and stories^13,33,34^. These results show that during verbal recall of a 50-minute movie, the neural patterns associated with individual scenes were reactivated in the absence of any external cues. For analysis of reinstatement at a finer temporal scale than the scene level, see Fig. S1.

**Pattern similarity between participants**. The preceding results established that freely spoken recall of an audio-visual narrative could elicit reinstatement of stimulus-induced activity patterns in an array of high-level cortical regions, including those that are typically observed during episodic memory retrieval^32^. Having mapped movie-vs.-recall correlations *within* individual participants, we next searched for correlations *between* participants during both movie and recall.

Previous studies have shown that viewing the same movie, or listening to the same story, can induce strong between-participant similarity in the timecourses of brain activity in many different regions^9,13,14,35^. Following this logic, we first examined similarities between participants during exposure to the same stimulus (the movie), but in the spatial domain rather than the temporal domain. Temporal and spatial aspects of neural activity, while not redundant, are certainly related in fMRI-recorded signals, and thus we expected to find similar areas as those previously observed for temporal similarity. Indeed, scene-specific spatial patterns of activity were highly similar across participants during movie-viewing in areas spanning the cortical hierarchy, from low-level sensory areas to higher-level association areas (Fig. S2). These results also echo prior studies that have effectively used cross-participant pattern analysis during shared perceptual stimulation in simpler paradigms^36–41^.

Next, we compared scene-specific stimulus-induced patterns (elicited during the movie) and scene-specific recollection patterns (elicited during recall) between participants. The analysis was identical to the reinstatement analysis described above (Fig. 2B), but performed *between* participants rather than *within* participant. For each participant, the recollection pattern for each scene was compared to the stimulus-induced pattern from that movie scene averaged across the remaining participants (Fig. 2D).

The searchlight analysis revealed extensive movie-vs.-recall correlations between participants, i.e., brain regions that had significantly similar scene-specific spatial patterns of activity between movie-viewing and spoken recall between participants (Fig. 2E-F). These results indicate that in many of the areas that exhibited movie-vs.-recall reinstatement effects within-individual, neural patterns elicited during spoken recollection in a given individual were similar to stimulus-induced neural patterns (elicited during the movie) in other individuals.

**Shared spatial patterns between participants during recall**. The preceding results showed that scene-specific neural patterns were shared across brains 1) during movie viewing, when all participants viewed the same stimulus (Fig. S2), and 2) when one participant’s recollection pattern was compared to other participants’ movie-induced brain patterns (Fig. 2D-F). Together, these results suggest that between-participants similarities might also be present during recollection, despite the fact that during the recall session no stimulus was presented and participants’ behavior differed drastically from each other (i.e., each participant described each movie scene in their own words and for different lengths of time). Thus, we next examined between-participants pattern similarity during the spoken recall session.

As before, brain data within each recall scene were averaged across time in each participant, resulting in one pattern of brain activity for each scene (“recollection pattern”). The recollection pattern from each scene for a given participant was compared directly to the recollection pattern for the same scene averaged across the remaining participants (Fig. 3A). Similarity was calculated using Pearson correlation. The analysis was performed in a searchlight across the brain volume and statistical significance, comparing the neural pattern similarity between matching scenes against that of non-matching scenes, was evaluated in the same manner as above (FDR corrected at q < 0.05, two-tailed).

**Figure 3.**
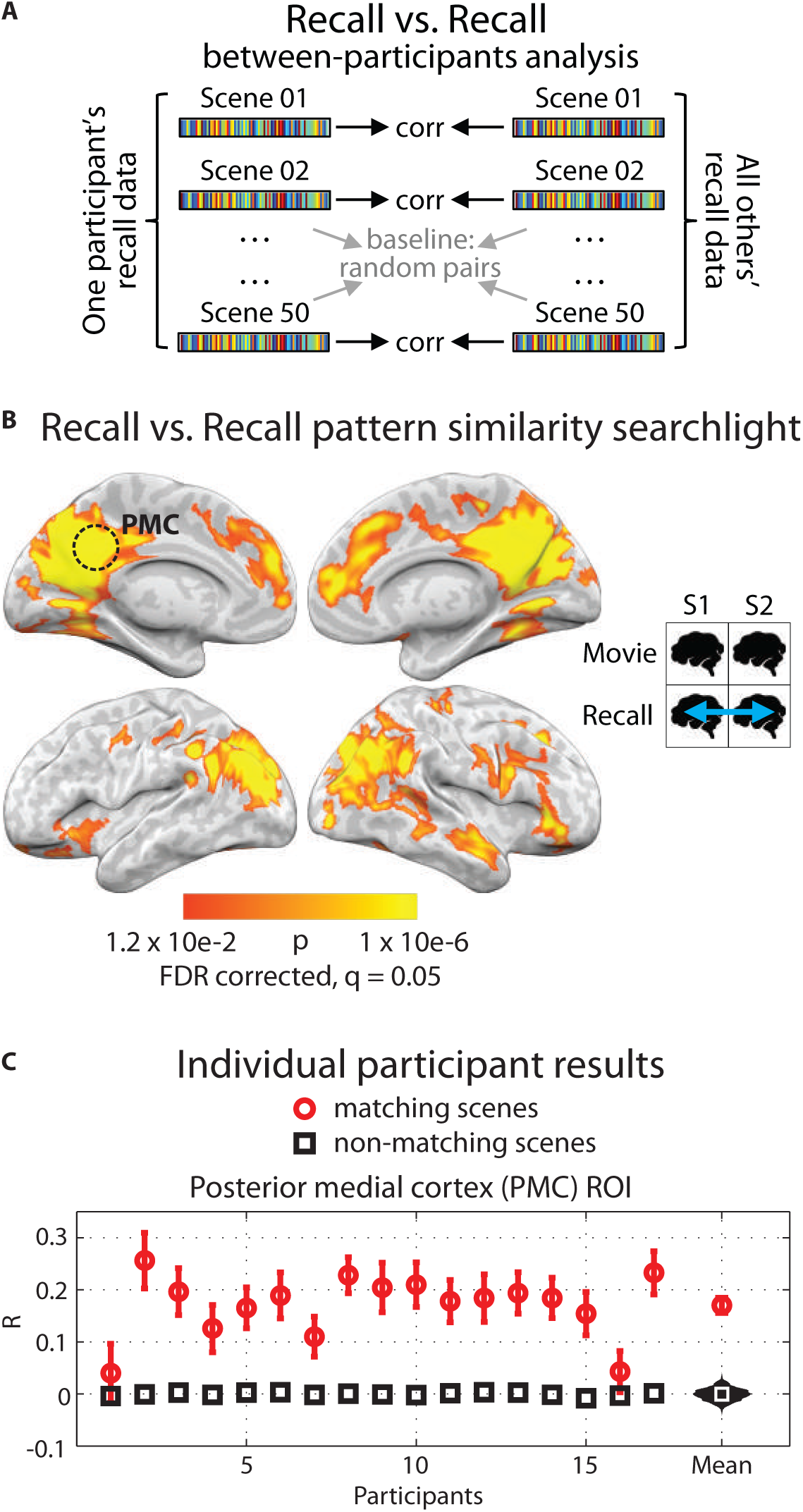
Between-participants pattern similarity during spoken recall. **A)** Schematic for between-participants recall-vs.-recall analysis. BOLD data from the recall sessions were divided into matching scenes, then averaged across time within each voxel, resulting in one vector of voxel values for each recalled scene. Next, correlations were computed between every matching pair of recalled scenes. Statistical significance was determined by shuffling scene labels (i.e., baseline correlations were calculated from non-matching movie/recall scene pairs) to generate a null distribution of the participant average. **B)** Searchlight map showing regions where significant recall-vs.-recall similarity was observed; FDR correction at q = 0.05, p = 0.012. Searchlight was a 5×5×5 voxel cube. **C)** Recall-vs.recall correlation values for all 17 participants in independently-defined PMC (posterior medial cortex). Red circles show average correlation of matching scenes and error bars represent standard error across scenes; black squares show average of the null distribution. At far right, the red circle shows the true participant average and error bars represent standard error across participants; black histogram shows the null distribution of the participant average; white square shows mean of the null distribution. See also Figure S3.

The searchlight analysis revealed a large set of brain regions that had significantly similar scene-specific patterns of activity between participants during spoken recall of shared experiences (Fig. 3B), including high-order cortical regions throughout the DMN as well as category-selective high-level visual areas, but not low-level sensory areas (see Fig. S3 for overlap with visual and auditory areas). Individual participant correlation values for independently-defined posterior medial cortex (PMC) are shown in Fig. 3C. These between-participants similarities were elicited despite the fact that no stimulus was present during recall, and individuals’ behavior – the compression factor of recollection and the words chosen by each person to describe each event the movie – varied dramatically. The direct spatial correspondence of event-specific patterns between individuals reveals the existence of a spatial organization to the neural representations that is common across brains.

**Classification accuracy**. How discriminable were the neural patterns for individual scenes? To address this question we performed a multi-voxel classification analysis^42^. Participants were randomly assigned to one of two groups (N=8 and N=9), and an average was calculated within each group for the PMC ROI (481 voxels). Pairwise correlations were calculated between the two group means for all 50 movie scenes. For any given scene (e.g., scene 1, group 1), the classification was labeled “correct” if the correlation with the matching scene in the other group (e.g., scene 1, group 2) was higher than the correlation with any other scene (e.g., scenes 2–50, group 2). Accuracy was then calculated as the proportion of scenes correctly identified out of 50. Classification rank was calculated for each scene (i.e., the rank of the matching scene correlation in the other group among all 50 scene correlations). The entire procedure was repeated using every possible combination of two groups sized N=8 and N=9 (Fig. 4A, green markers) and averaged. Overall classification accuracy was 38.4%, p < 0.001 (Fig. 4A, black bar; chance level [2.0%] plotted in red); classification rank for individual scenes was significantly above chance for 49 of 50 scenes, p < 0.001 (Fig. 4B, mean rank across scenes 4.5 out of 50). For both Fig. 4A and 4B, statistical significance was evaluated using a permutation analysis in which scene labels were randomized across the two groups, and corrected for multiple comparisons across the 50 scenes using FDR (q < 0.05, two-tailed).

**Figure 4.**
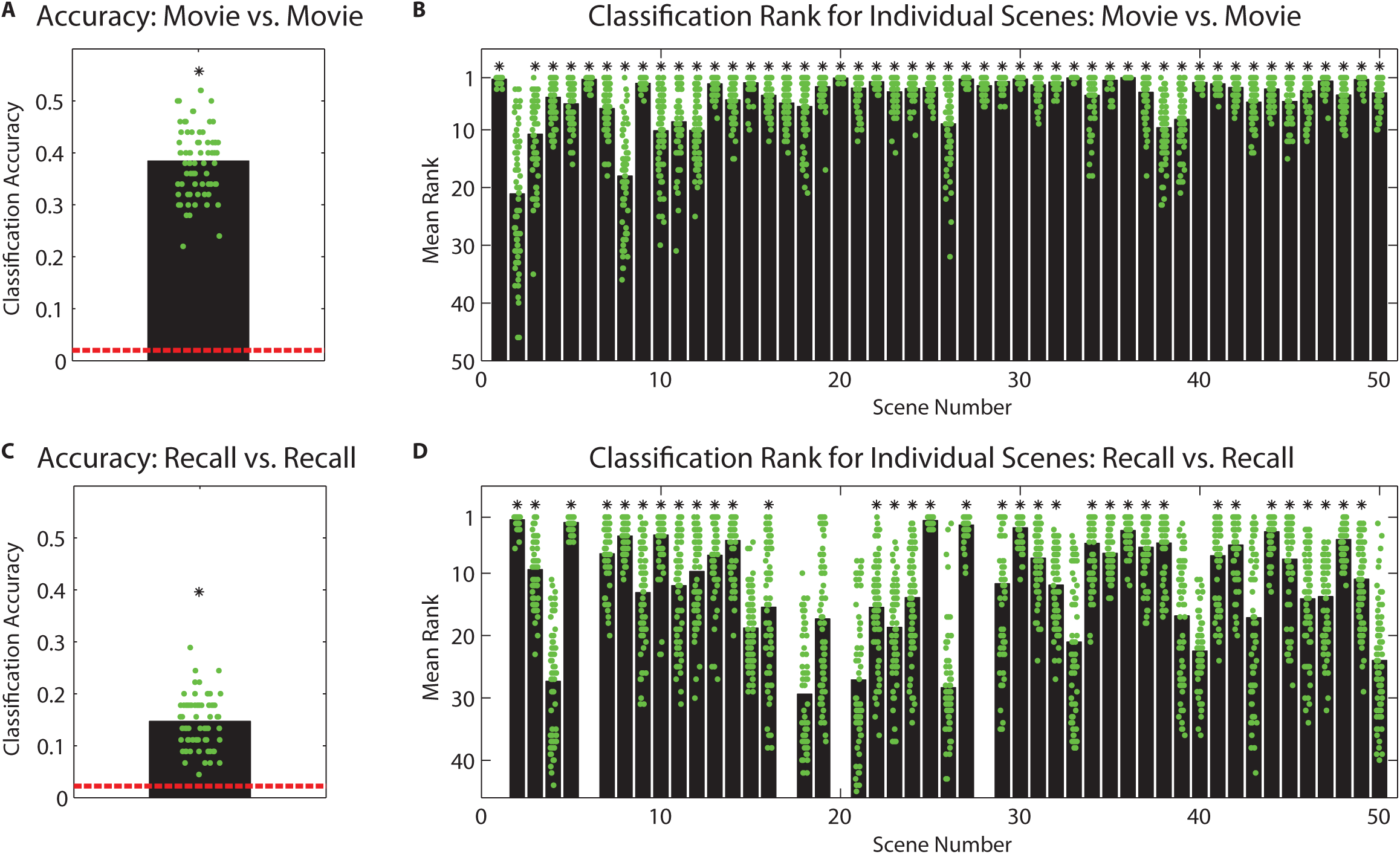
Classification accuracy. **A)** Classification of movie scenes between brains. Participants were randomly assigned to one of two groups (N=8 and N=9), an average was calculated within each group, and data were extracted for the PMC ROI. Pairwise correlations were calculated between the two group means for all 50 movie scenes. Accuracy was then calculated as the proportion of scenes correctly identified out of 50. The entire procedure was repeated using every possible combination of two groups sized N=8 and N=9 (green markers), and an overall average calculated (38.4%, p < 0.001, black bar; chance level [2.0%] plotted in red). **B)** Classification rank for individual movie scenes (i.e., the rank of the matching scene correlation in the other group among all 50 scene correlations). Green markers show the results from each combination of two groups sized N=8 and N=9; black bars show the average over all group combinations, 4.5 on average. (* indicates p < 0.001, FDR corrected.) **C)** Classification of recalled scenes between brains. Same analysis as in (A) except that data were extant for 45 scenes. Overall classification accuracy was 18.0% (p < 0.001, black bar, chance level 2.2%). **D)** Classification rank for individual recalled scenes, 11.5 on average. (* indicates p < 0.001, FDR corrected.)

We next computed the same classification analyses using the data from recall. The analyses were identical to those above with the exception that data were not extant for all 50 scenes for every participant, due to participants recalling 34.4 scenes on average. Thus, group average patterns for each scene were calculated by averaging over the extant data; for 45 scenes there were data available for at least one participant in each group, considering all possible combinations of participants into groups of N=8 and N=9. Overall classification accuracy was 18.0% (Fig. 4C; chance level 2.2%). Classification rank for individual scenes was significantly above chance for 34 of 45 possible scenes, p < 0.001 (Fig. 4D; mean rank across scenes 11.5 out of 45). For both Fig. 4C and 4D, statistical significance was evaluated using a permutation analysis in which scene labels were randomized across the two groups, and corrected for multiple comparisons across the 45 scenes using FDR (q < 0.05, two-tailed). See also Fig. S4.

**Transformation of neural patterns from perception to recollection**. A key question of the experiment was how neural representations change between perception (the movie) and memory (recollection). To address this question, we conceptualized the transformation between percept-based neural patterns and recollection patterns in the following way:

For each participant, the percept-based neural patterns for a given scene are expressed as a common underlying pattern, plus idiosyncratic responses. Each of these patterns is then altered in some manner in order to produce the recollection pattern. If representations are changed in a unique way within each person’s brain, then each person’s percept-based (movie) pattern is altered by adding (or multiplying by) a “transformation” pattern that is uncorrelated with the “transformation” patterns of other people. In this scenario, recollection patterns for a given scene necessarily become more dissimilar to the recollection patterns of other people as compared to the pattern elicited during the original movie scene (Fig. 5A, left).

**Figure 5.**
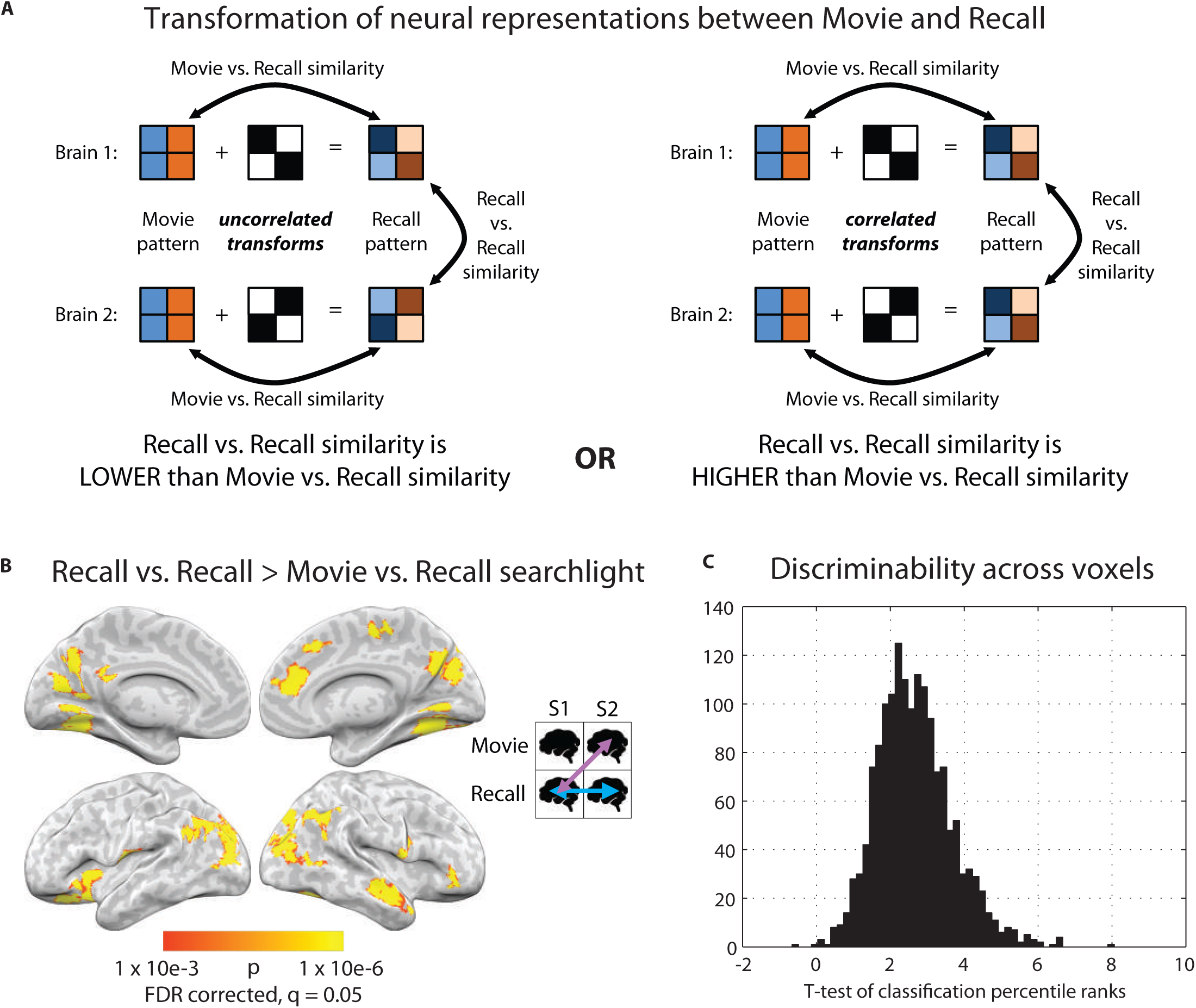
Transformation of neural patterns from perception to recollection. **A)** Schematic showing how neural activity patterns during a movie scene are transformed into activity patterns at recall. For each brain, the percept-based neural patterns for a given scene are expressed as a common underlying pattern. Each of these Movie patterns is then altered in some manner to produce the Recall pattern. Left panel: If patterns are changed in a unique way within each person's brain, then each person's percept-based (movie) pattern is altered by adding a “transformation” pattern that is uncorrelated with the “transformation” patterns of other people. In this scenario, Recall patterns necessarily become more dissimilar to the Recall patterns of other people than to the Movie pattern. Right panel: Alternatively, if a systematic change is ocurring across people, this corresponds to a scenario in which each Movie pattern is altered by adding a “transformation” pattern that is correlated with the “transformation” patterns of other people. Thus, Recall patterns for a given scene may become more similar to the Recall patterns of other people than to the Movie pattern (Fig. 5A, right). **B)** Searchlight map showing regions where recall-vs.-recall similarity was significantly greater than between-participants movie-vs.-recall similarity, i.e., where the map from Fig. 3B was stronger than the map from Fig. 2E. Significance was calculated using a bootstrap analysis wherein individual participant values were randomly swapped between conditions. P-values were corrected for multiple comparisons across the entire brain using FDR (q < 0.05). The analysis revealed an array of regions, including posterior parahippocampal cortex, right superior temporal pole, posterior medial cortex, right medial prefrontal cortex, and angular gyrus, in which neural representations changed in a a systematic way across individuals between perception and recollection. **C)** We tested whether each participant's individual scene recollection patterns could be classified better using 1) the movie data from other participants, or 2) the recall data from other participants. A t-test of classification rank was performed between these two sets of values at each voxel underlying the searchlight cubes in (B). Classification rank was higher when using the recall data as opposed to the movie data in 99.9% of such searchlights. Histogram of t-values is plotted.

Alternatively, if a systematic change is ocurring across people, this corresponds to a scenario in which each percept-based (movie) pattern for a given scene is altered by adding (or multiplying by) a “transformation” pattern that is *correlated* with the “transformation” patterns of other people. In this case, recollection patterns for a given scene may become *more similar* to the recollection patterns of other people than to the pattern elicited during the original movie scene (Fig. 5A, right).

Thus, we looked for brain regions in which, for individual scenes, recollection activity patterns were more similar to recollection patterns in other individuals than they were to movie patterns. In order to ensure a balanced contrast, we compared the between-participants recall-vs.-recall values to the between-participants movie-vs.-recall values (rather than to within-participant movie-vs.-recall). Statistical significance of the difference was calculated using a permutation test and FDR-corrected across all voxels in the brain. The analysis revealed an array of regions, including posterior parahippocampal cortex, right superior temporal pole, posterior medial cortex, right medial prefrontal cortex, and angular gyrus (Fig. 5B), in which neural representations changed in a systematic way across individuals between perception and recollection. If representations were modified between encoding and recall in a significantly idiosyncratic manner across individuals, we would expect the exact opposite result: lower recall-vs.-recall similarity than movie-vs.-recall similarity. No regions in the brain showed such a pattern.

A possible concern was that the greater similarity for recall-vs.-recall relative to movie-vs.-recall might arise simply from the fact that all recall data were collected during speech, while there was no speech during the movie. In other words, that the greater similarity might be due merely to the common physical activity (i.e., speech) across recalled events. To test this concern, we examined the *discriminability* of individual scenes in more detail. While our measure of pattern correlation is already a scene-specific measure, in that matching scenes are shown to be more strongly correlated than nonmatching scenes, the discriminability of scenes is tested more rigorously by calculating classification success for individual scenes. Within the regions shown in Fig. 5B (1568 voxels), we asked whether each participant’s individual scene recollection patterns could be classified better using a) the movie data from other participants, or b) the recall data from other participants. Classification rank was calculated (the rank of the matching scene correlation among all of the recalled scene correlations); this was averaged across scenes to produce one value per participant for (a) and one for (b). The analysis was performed for every searchlight underlying the voxels shown in Fig. 5A. Classification rank was higher when using the recall data as opposed to the movie data in 99.9% of such searchlights (see Fig. 5C for the distribution of t-values resulting from a test between recall-vs.-recall and movie-vs.-recall).

Together, these results show that, in a subset of DMN regions, neural representations of each event were modified between perception and recall in a systematic manner across individuals.

**Subsequent memory**. To examine how neural activity during movie and recall might be related to memorability, we divided scenes into remembered and forgotten for each participant. For each scene we calculated how many participants had successfully recalled that scene, and examined recollection patterns for each scene in PMC. Fig. 6A shows that, for any given scene, the strength of neural recall-vs.-recall correlation was significantly related to how likely that scene was to be remembered (R = 0.52, p < 0.001). In other words, the more that a given movie scene gave rise to shared recollection neural patterns across participants, the more likely that scene was to be remembered. A similar result was found when using all voxels in the default mode network (R = 0.38, p < 0.01). A control analysis showed that between-participants movie-vs.-recall correlation was not predictive of the likelihood of recall (R = 0.00, p > 0.9, Fig. 6B). See Fig. 6C-D for additional control analyses.

**Figure 6.**
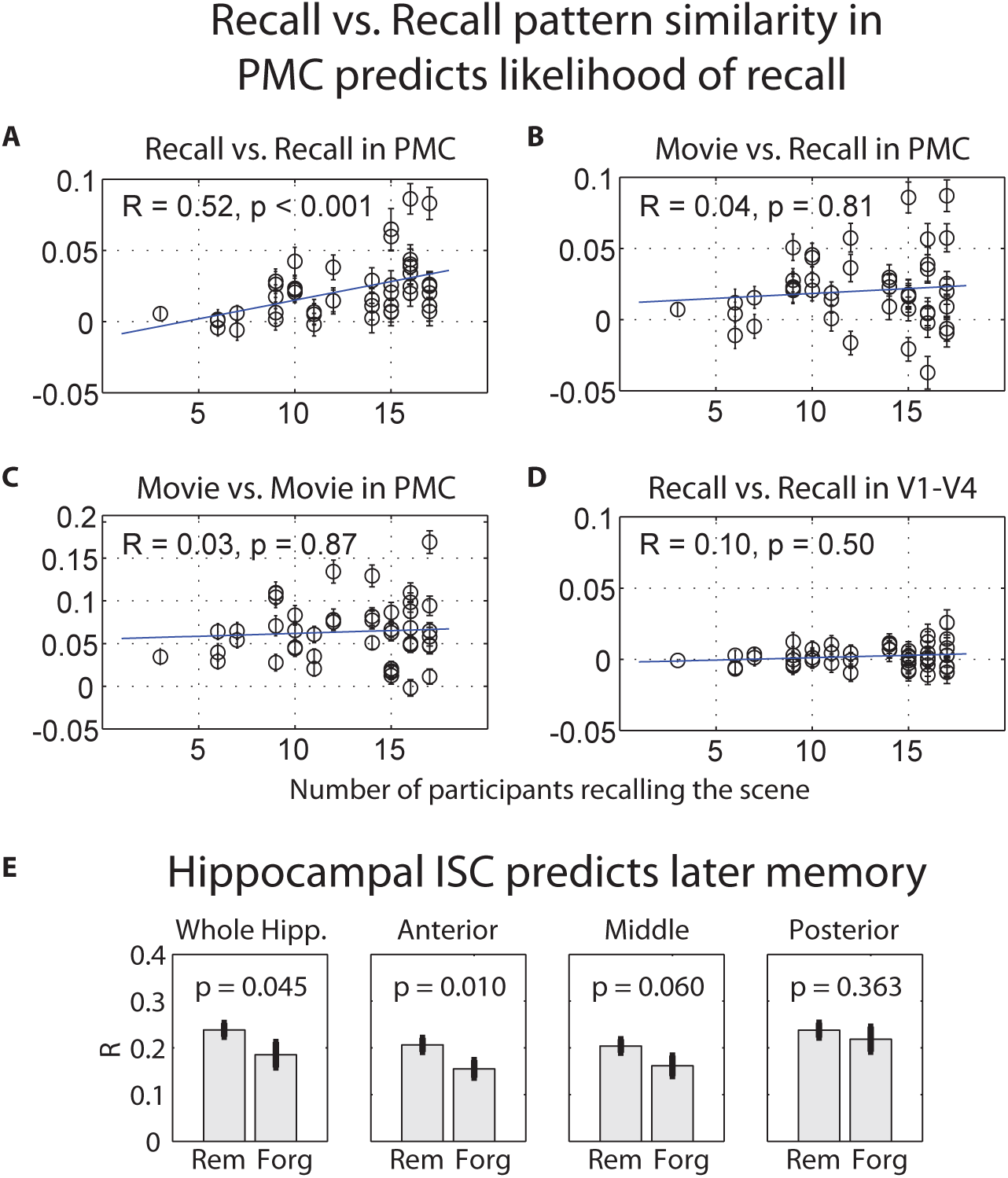
Subsequent memory analyses. **A)** The number of participants who successfully recalled each scene was calculated, and patterns for each scene were examined in the PMC ROI. Pattern similarity between participants was calculated in a pairwise manner for each scene during recall. The average similarity between participants for each scene during recall was significantly related to how likely that scene was to be recalled (Spearman rank correlation: R = 0.52, p < 0.001). In other words, the more that a given movie scene gave rise to similar recall patterns between participants, the more likely that scene was to be remembered. **B)** A control analysis showing that between-participants pattern similarity between movie and recall was not predictive of the likelihood of recall in PMC (R = 0.04, p > 0.8). **C)** A control analysis showing that between-participants pattern similarity during movie-viewing was not predictive of the likelihood of recall in PMC (R = 0.03, p > 0.8). **D)** A control analysis showing that between-participants pattern similarity during recall in early visual areas V1-V4 was not predictive of the likelihood of recall (R = 0.10, p = 0.5, same ROI as Fig. S3). **E)** Hippocampal inter-subject correlation (ISC) over time was calculated for each movie scene in each participant and the scenes were binned by whether they were later remembered or forgotten. Using a whole hippocampus ROI, ISC was significantly greater for remembered scenes than forgotten scenes (left panel; 2-tailed paired t-test across participants, p = 0.045). The same analysis is shown for the hippocampus ROI split into anterior, middle, and posterior sections (second, third, and fourth panels from the left). A repeated-measures ANOVA with region (anterior, middle, posterior) and memory (remembered, forgotten) as factors revealed significant main effects of region F(2,32) = 12.02, p < 0.0005 and of memory F(1,16) = 4.98, p = 0.04, but not a significant region x memory interaction F(2,32) = 1.69, p = 0.2.

As the task depended on episodic memory, we also examined hippocampal contributions. During movie viewing, we calculated the correlation between a given participant’s hippocampal timecourse and the average hippocampal timecourse of all other participants, for individual scenes (i.e., the inter-subject correlation (ISC) for each scene^9^). This hippocampal ISC was significantly predictive of which scenes would later be recalled (remembered vs. forgotten: t = 2.17, p = 0.045; Fig. 6E), complementing previous results linking ISC in parahippocampal cortex to later recognition memory^43^.

**Visualization of BOLD activity in individual scenes**. In order to visualize the underlying signal, we randomly split the movie-viewing data into two independent groups of equal size (N=8 each) and averaged BOLD values at every voxel across participants within each group. An average was made in the same manner for the recall data using the same two groups of eight. These group mean images were then averaged across timepoints and within scene, creating one brain image per group per scene. For movie data see Fig. 7A (leftmost 2 panels) and 7B-C; for recall see Fig. 7A (rightmost 2 panels) and 7D-E. This averaging procedure reveals the component of the BOLD signal that is shared across brains, i.e., if a similar activity pattern can be observed between the two independent groups for an individual scene, it indicates a common neural response across participants. Visual inspection of these images suggests replication across groups for individual scenes, and differentiation of the shared signal between scenes, as quantified in the classification analysis above (see Fig. 4). See also Fig. S4 for individual scene correlation values.

**Figure 7.**
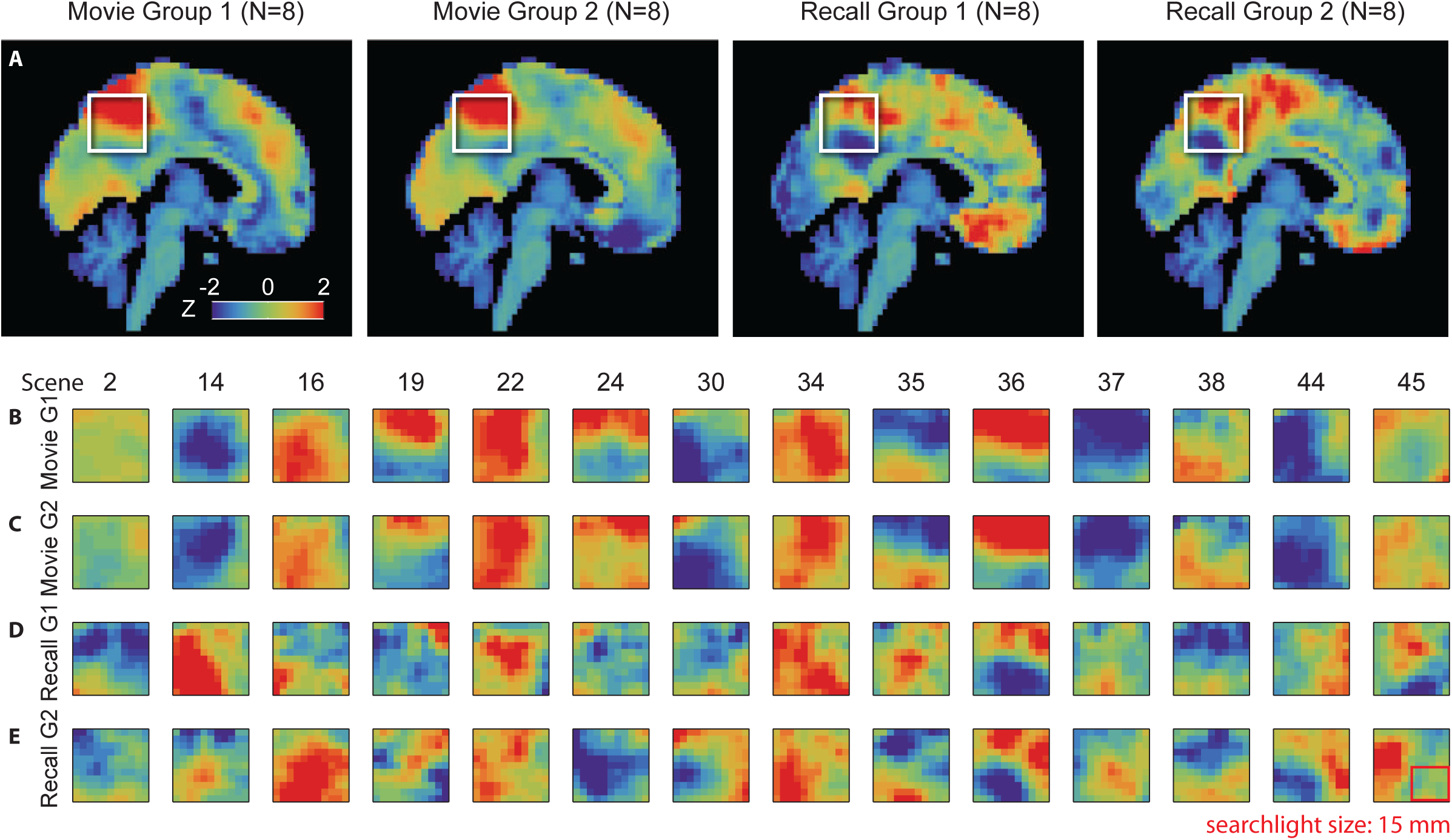
Scene-level pattern similarity between individuals. Visualization of the signal underlying pattern similarity between individuals, for fourteen scenes that were recalled by all sixteen of the participants in these groups, are presented in [B-E]. See Figure S4 for correlation values for all scenes. **A)** In order to visualize the underlying signal, we randomly split the movie-viewing data into two independent groups of equal size (N=8 each) and averaged BOLD values across participants within each group. An average was made in the same manner for the recall data using the same two groups of eight. These group mean images were then averaged across timepoints and within scene, exactly as in the prior analyses, creating one brain image per group per scene. Sagittal view of these average brains during one representative scene (36) of the movie is shown for each group. Average activity in a posterior-medial area (white box in [A]) on the same slice for the fourteen different scenes for Movie Group 1 **(B)**, Movie Group 2 **(C)**, Recall Group 1 **(D)**, and Recall Group 2 **(E)**. Searchlight size shown as a red outline.

**Reinstatement in individual participants vs. between participants**. While our between-participant analyses explored the common, shared component of memory representations, the question arises as to whether the neural patterns also contain information reflecting more fine-grained individual differences in memory representations (e.g.,^44^). If so, one would expect within-participant movie-vs.-recall similarity (reinstatement) to be stronger than between-participant movie-vs.-recall similarity. A simple comparison of within-participant movie-vs.-recall pattern similarity (Fig. 2B-C) to between-participant movie-vs.-recall pattern similarity (Fig. 2E-F) does not suffice, due to anatomical registration being better within-participant than between-participants.

In order to mitigate the anatomical misalignment bias, we performed a second-order similarity analysis: correlation of representational dissimilarity matrices (RDMs)^30,44^ within and between participants. Each RDM was composed of the pairwise correlations of patterns for individual scenes in the movie (“movie-RDM”) and during recall (“recall-RDM”) calculated within participants. The movie-RDMs map the relationships between all movie scenes, and the recall-RDMs map the relationships between all recalled scenes. Because the RDMs were always calculated within-brain, we were able to assess the similarity between representational structures within and between participants (by comparing the movie-RDMs to the recall-RDMs within-participant and between-participants [see Fig. 8A-B insets]) in a manner less susceptible to potential anatomical misalignment between participants.

**Figure 8.**
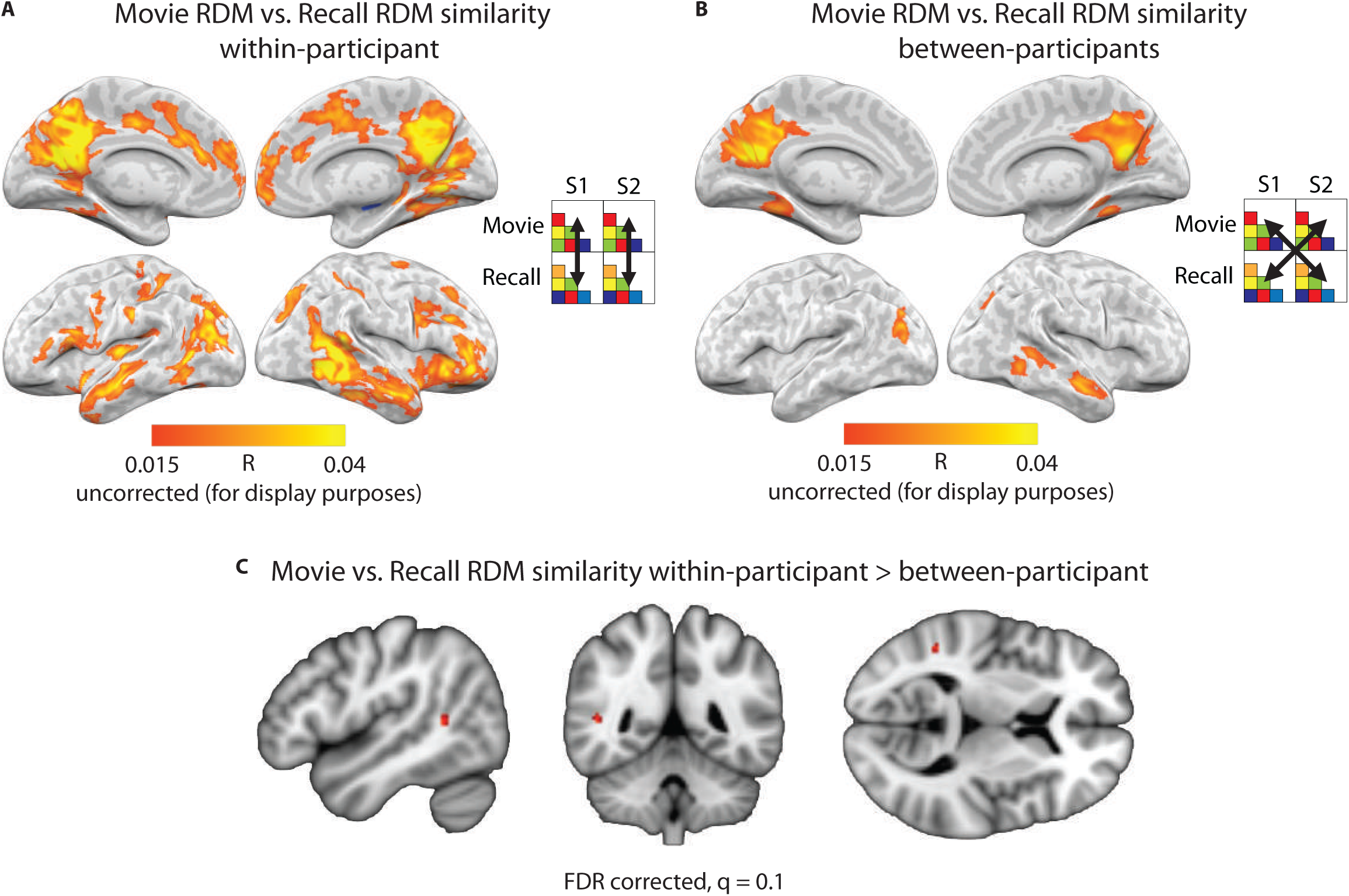
Reinstatement in individual participants vs. between participants. **A**) Searchlight analysis showing similarity of representational dissimilarity matrices (RDMs) within-participant across the brain. Each RDM was composed of the pairwise correlations of patterns for individual scenes in the movie (“movie-RDM”) and separately during recall (“recall-RDM”). Each participant's movie-RDM was then compared to his or her own recall-RDM (i.e., within-participant) using Pearson correlation. The average searchlight map across 17 participants is displayed. **B)** Searchlight analysis showing movie-RDM vs. recall-RDM correlations between participants. The average searchlight map across 272 pairwise combinations of participants is displayed. **C)** The difference was computed between the within-participant and between-participant maps. Statistical significance of the difference was evaluated using a permutation analysis and FDR corrected (q < 0.05, two-tailed). A cluster of two voxels located in the temporo-parietal junction survived correction (map shown at q < G.1G for visualization purposes, 5-voxel cluster).

We calculated movie-RDM vs. recall-RDM correlations, within-participant, in a searchlight analysis across the brain volume (Fig. 8A). The same analysis was performed for all pairs of participants (Fig. 8B). Of critical interest was the difference between the within-participant comparison and the between-participant comparison. Statistical significance of the difference was evaluated using a permutation analysis that randomly swapped condition labels for within- and between-participant RDM correlation values, and FDR corrected across all voxels in the brain (q < 0.05, two-tailed). This analysis revealed a single cluster located in the right temporoparietal junction (MNI coordinates: 48, −48, 9; Fig. 8C) for which within-participant movie-RDM vs. recall-RDM correlation was significantly greater than between-participants movie-RDM vs. recall-RDM correlation, i.e., in which individual-unique aspects of neural patterns contributed to reinstatement strength above and beyond the shared representation.

## DISCUSSION

In this study, we found that patterns of brain activity are reactivated as individuals recall an event. Crucially, we also found that these patterns are similar across individuals remembering the same event. This shared activity was observed during free spoken recall as participants reported the contents of their memories (a movie they had watched) in their own words, in the absence of any sensory cues or experimental intervention. A large set of high-order multimodal cortical areas was implicated, including posterior medial cortex, medial prefrontal cortex, parahippocampal cortex, and posterior parietal cortex, as well as high-level visual areas such as face- and scene-selective regions of ventral temporal cortex, and attentional regions in intraparietal sulcus. The direct spatial correspondence of event-specific patterns between individuals reveals the existence of a spatial organization to the neural representations that is common across brains. In a subset of regions, individual participants’ recollection patterns were more similar to the recollection patterns of others than to patterns elicited during the movie itself, suggesting that neural representations of each event were modified between perception and recall in a structured manner across individuals. Furthermore, the degree to which recollection patterns were shared across participants predicted the memorability of individual movie scenes. Overall, these findings show that memory representations have a common spatial organization in the brain, and that perceptual experience is transformed into memory in a systematic manner across different individuals, even as people speak freely in their own words about past events.

The brain areas in which we observed shared representations during recall overlap strongly with the “default mode network” (DMN)^8^. The DMN has been implicated in a broad range of complex cognitive functions, including scene and situation model construction, episodic memory, and internally focused thought^31,32,45–48^. Multiple studies have shown that during processing of real-life stimuli such as movies and stories, activity timecourses in these regions are synchronized across individuals and locked to high-level aspects of the stimulus, but not to low-level sensory features. For example, these regions evince the same narrative-locked coherent dynamics whether a given narrative is presented in spoken or written form^49,50^, and whether it is presented in English or Russian^14^. Dynamics in these regions are modulated according to the perspective of the perceiver^16^, but when comprehension of the narrative is disrupted (while keeping low-level sensory features unchanged), neural activity becomes incoherent across participants^13,14,51^. Together, these results suggest that representations in the DMN track high-level information structure (e.g., narrative or situational elements^46^) in the input. The current study extends prior findings by demonstrating that these high-level DMN representations formed during encoding can be reinstated at will from memory, without the need for any guiding stimulus. That is, the shared neural responses among individuals while experiencing the same events later give rise to shared neural responses during recollection, even when each person describes the past in his or her own words.

A memory is not a perfect replica of the original experience; perceptual representations undergo modification in the brain prior to recollection that may increase the usefulness of the memory, e.g., by emphasizing certain aspects of the percept and discarding others. What laws govern how neural representations change between perception and memory? We examined whether the transformation of neural patterns from percept to memory was idiosyncratic as opposed to systematic across people. We reasoned that if each person’s perceptual experience is transformed into memory in a unique manner, neural patterns should become *more dissimilar* across individuals recalling the same event than to the original percept-based pattern. Alternatively, if perceptual experience is transformed into memory in a structured way, then reactivated patterns might change systematically to become *more similar* across individuals than to the original percept-based pattern. Recall-vs.-recall similarity across participants during recollection was stronger than movie-vs.-recall similarity in several brain regions, including posterior parahippocampal cortex, superior temporal pole, posterior medial cortex, and medial prefrontal cortex (Fig. 5B). Not only the similarity, but also the discriminability of events was increased during recall, indicating that the greater similarity was not due to a common factor (i.e., speech) across recalled events. Interestingly, we found that scenes which were the most neurally similar between individuals during recollection were also the most likely to be recalled (Fig. 6A). A possible interpretation of these findings is that participants shared familiar notions of how certain events are structured (for example, what elements are typically present in a car chase scene in an action movie), and that these existing schemas guided encoding and/or recall of new events. Such forms of shared knowledge might improve later memory by allowing participants to think of schema-consistent items or events, essentially providing self-generated memory cues^52^.

How refined was the spatial alignment of recollection patterns across brains? The alignment had to be robust enough to overcome imperfect registration (due to individual variation in brain anatomy) across the brains of different participants. Our data indicate that the pattern alignment was strong enough to survive spatial transformation of brain data to a standard anatomical space, even when using a small (5^3^ voxels) searchlight size. Recently, it was argued that relatively coarse organized patterns (e.g., small eccentricity-related biases toward horizontal or vertical orientations within primary visual cortex) can underlie spatial pattern correlations in the brain^53^. Our findings suggest that memory representations, similar to sensory representations^54^, are organized in a functional architecture that is shared across participants. Importantly, while the current results reveal a relatively coarse spatial structure that is shared across people (Fig. 7), they do not preclude the existence of a finer spatial structure in the neural signal that may be captured when comparisons are made within-participant^44,55^ or by using more sensitive methods such as hyperalignment^38^. When we calculated movie-RDM vs. recall-RDM second-order correlations within and between brains, we found only a single region in the temporoparietal junction in which individual-unique aspects of neural patterns contributed reliably to reinstatement strength above and beyond the shared representation. The fact that this region comprises a relatively small proportion of the total cortical area in which we identified shared representations suggests that a substantial portion of the movie-vs.-recall pattern similarity was captured in the between-brain comparison. Using a similar second-order correlation approach, Charest et al. (2014) found individual-unique neural responses in inferior temporal cortex. However, note that there were numerous differences between our study and the Charest et al. study, in stimulus content (Charest et al. used objects selected to have personal significance to the participants), paradigm, and regions of interest; further work is needed to understand the factors that influence the balance of idiosyncratic and shared signals between brains.

Could the scene-specific patterns observed during movie and recall be explained by varying levels of arousal across the scenes? One might imagine that univariate activity, scaling with arousal, could provide some information relevant to scene decoding. For example, if response patterns to low-arousal scenes differed systematically from high-arousal scenes, and half of the 50 scenes were low-arousal, then classification accuracy could reach 4% by this factor alone (1/50 * 2 differentiable levels of arousal). In Fig. 4 we show that classification of individual movie scenes between brains was 38.4% (chance level 2.0%), and classification of individual recalled scenes between brains was 18.0% (chance level 2.9%) in PMC. In order to reach these levels of performance under the framework described above, there would have to be 19 differentiable levels of arousal during the movie and 9 differentiable levels during recall. Thus, while it is likely that arousal plays some role in neural differentiation of scenes, it seems insufficient to explain the full pattern of results.

To what extent did spoken recollection in this study engage visual imagery? We observed extraordinarily rich recollection behavior (Fig. 1B, Table S2), in which participants managed to recount, in largely correct order, the details of most of the movie scenes; this suggested that participants were able to mentally replay virtually the entire movie, despite the absence of any external cues. However, movie-vs.-recall reinstatement effects were not found in low level visual areas, but instead were located in high level visual areas (Fig. S3), and extensively in higher order brain regions outside of the visual system (Fig. 2). Our observation of reinstatement in high level visual areas is compatible with studies showing reinstatement in these regions during cued visual imagery^25,27,28,56^. The lack of reinstatement effects in low-level areas may be due to the natural tendency of most participants to focus on the episodic narrative (the plot) when recounting the movie, rather than on fine visual details. It has been suggested that the requirement to note high-resolution details is a key factor in eliciting activity in early visual cortex during visual imagery^57^. Thus, our findings do not conflict with studies showing that activity patterns in early visual cortex can be used to decode a simple image held in mind during a delay, in tasks that required vivid imagery of low-level visual features^58,59^.

Together, these results show that a common spatial organization for memory representations exists in high-level cortical areas (e.g., the DMN), where information is largely abstracted beyond sensory constraints; and that perceptual experience is transformed before recall in a systematic manner across people, a process that may benefit memory. These observations were made as individuals engaged in natural and unguided spoken recollection, testifying to the robustness and ecological validity of the phenomena. Future work may explore whether these shared representations underlie our ability to share memories with others^60^, and how they might contribute to a community’s collective memory ^2–7^

## Acknowledgments

We thank C. Honey, M. Aly, C. Baldassano, M. Arcaro, and E. Simony for scientific discussions and helpful comments on earlier versions of the manuscript; J. Edgren for help with transcription; M. Arcaro for advice regarding visual area topography; and other members of the Hasson and Norman labs for their comments and support. This work was supported by The National Institutes of Health (R01-MH094480, 2T32MH065214-11).

## Methods

### Participants

Twenty-two participants were recruited from the Princeton community (12 male, 10 female, ages 18–26, mean age = 20.8). All participants were right-handed native English speakers, reported normal or corrected-to-normal vision, and had not watched any episodes of *Sherlock*^21^ prior to the experiment. All participants provided informed written consent prior to the start of the study in accordance with experimental procedures approved by the Princeton University Institutional Review Board. The study was approximately two hours long and participants received $20 per hour as compensation for their time. Data from five out of the twenty-two participants were discarded due to excessive head motion (greater than one voxel; 2 participants), or because recall was shorter than 10 minutes (2 participants), or for falling asleep during the movie (1 participant).

### Stimuli

The audio-visual movie stimulus was a 48-minute segment of the BBC television series *Sherlock*^21^, taken from the beginning of the first episode of the series (a full episode is 90 minutes). The stimulus was further divided into two segments (23 and 25 minutes long); this was done to reduce the length of each individual runs, as longer runs might be more prone to technical problems (e.g., scanner overheating). At the beginning of each of the two movie segments, a 30-second audiovisual cartoon was prepended *(Let’s All Go to the Lobby*^61^) that was unrelated to the *Sherlock* movie.

### Experimental Procedures

Participants were told that they would be watching the British television crime drama series *Sherlock*^21^ in the fMRI scanner. They were given minimal instructions: to attend to the audiovisual movie, e.g., “watch it as you would normally watch a television show that you are interested in,” and that afterward they would be asked to verbally describe what they had watched. Participants then viewed the 50-minute movie in the scanner. The scanning (and stimulus) was divided into two consecutive runs of approximately equal duration.

The movie was projected using an LCD projector onto a rear-projection screen located in the magnet bore and viewed with an angled mirror. The Psychophysics Toolbox [http://psychtoolbox.org] for MATLAB was used to display the movie and to synchronize stimulus onset with MRI data acquisition. Audio was delivered via in-ear headphones. Eyetracking was conducted using the iView X MRI-LR system (Sensomotoric Instruments [SMI]). No behavioral responses were required from the participants during scanning, but the experimenter monitored participants’ alertness via the eyetracking camera. Any participants who appeared to fall asleep, as assessed by video monitoring, were excluded from further analyses.

At the start of the spoken recall session, which took place immediately after the end of the movie, participants were instructed to describe what they recalled of the movie in as much detail as they could, to try to recount events in the original order they were viewed in, and to speak for at least 10 minutes if possible but that longer was better. They were told that completeness and detail were more important than temporal order, and that if at any point they realized they had missed something, to return to it. Participants were then allowed to speak for as long as they wished, and verbally indicated when they were finished (e.g., “I’m done”). During this session they were presented with a static black screen with a central white dot (but were not asked to, and did not, fixate); there was no interaction between the participant and the experimenter until the scan ended. Functional brain images and audio were recorded during the session. Participants’ speech was recorded using a customized MR-compatible recording system (FOMRI II; Optoacoustics Ltd.).

### Behavioral Analysis

Timestamps were identified that separated the audiovisual movie into 48 “scenes”, following major shifts in the narrative (e.g., location, topic, and/or time). These timestamps were selected by an independent coder with no knowledge of the experimental design or results. The scenes ranged from 11 to 180 [s.d. 41.6] seconds long. Each scene was given a descriptive label (e.g., *Press conference)*. Together with the two identical cartoon segments, this resulted in 50 total scenes.

Transcripts were written of the audio recording of each participant’s spoken recall. Timestamps were then identified that separated each audio recording into the same 50 scenes that had been previously selected for the audiovisual stimulus. A scene was counted as “recalled” if the participant described any part of the scene. Scenes were counted as “out of order” if they were initially skipped and then described later. See Tables S1 and S2.

### fMRI Acquisition

MRI data were collected on a 3T full-body scanner (Siemens Skyra) with a 20-channel head coil. Functional images were acquired using a T2*-weighted echo planar imaging (EPI) pulse sequence (TR 1500 ms, TE 28 ms, flip angle 64, whole-brain coverage 27 slices of 4 mm thickness, in-plane resolution 3 × 3 mm^2^, FOV 192 × 192 mm^2^), with ascending interleaved acquisition. Anatomical images were acquired using a T1-weighted MPRAGE pulse sequence (0.89 mm^3^ resolution).

### fMRI Analysis

#### Preprocessing

Preprocessing was performed in FSL [http://fsl.fmrib.ox.ac.uk/fsl], including slice time correction, motion correction, linear detrending, high-pass filtering (140 s cutoff), and coregistration and affine transformation of the functional volumes to a template brain (MNI). Functional images were resampled to 3 mm isotropic voxels for all analyses. All calculations were performed in volume space. Projections onto a cortical surface for visualization were performed, as a final step, with NeuroElf (http://neuroelf.net).

Motion was minimized by instructing participants to remain very still while speaking, and stabilizing participants’ heads with foam padding. Artifacts generated by speech may introduce some noise, but cannot induce positive results, as our analyses depend on spatial correlations between sessions (movie vs. recall or recall vs. recall). Similar procedures regarding speech production during fMRI are described in previous publications from our group^62,63^.

#### ROI definition

An anatomical hippocampus ROI was defined based on the probabilistic Harvard-Oxford Subcortical Structural atlas^64^, and an ROI for posterior medial cortex (PMC) was taken from an atlas defined from resting-state connectivity^65^: specifically, the posterior medial cluster in the “dorsal default mode network” set (http://findlab.stanford.edu/functional_ROIs.html). A default mode network ROI was created by calculating the correlation between the PMC ROI and every other voxel in the brain (i.e., “functional connectivity”) during the movie for each subject, averaging the resulting maps across all subjects, and thresholding at R = 0.4.

#### Pattern similarity analyses

The brain data were transformed to standard MNI space. For each participant, data from movie-viewing and spoken recall were each divided into the same 50 scenes as defined for the behavioral analysis. BOLD data were averaged across timepoints within-scene, resulting in one pattern of brain activity for each scene: one “stimulus-induced pattern” elicited during each movie scene, and one “recollection pattern” elicited during spoken recall of each scene. Recollection patterns were only available for scenes that were successfully recalled, i.e., each participant possessed a different subset of recalled scenes. Each scene-level pattern could then be compared to any other scene-level pattern in any region (e.g., an ROI or a searchlight cube). All such comparisons were made using Pearson correlation.

For searchlight analyses^29^, pattern similarity was calculated in 5 × 5 × 5 voxel cubes (i.e., 15 × 15 × 15 mm cubes) centered on every voxel in the brain. Statistical significance was determined by shuffling scene labels to generate a null distribution of the average across participants, i.e., baseline correlations were calculated from non-matching scene pairs^30^. This procedure was performed for each searchlight cube, with one cube centered on each voxel in the brain; cubes with 50% or more of their volume outside the brain were discarded. The results were corrected for multiple comparisons across the entire brain using FDR (q < 0.05). Importantly, the nature of this analysis ensures that the discovered patterns are content-specific at the scene level, as the correlation between neural patterns during matching scenes must on average exceed correlations between non-matching scenes in order to be considered statistically significant.

Four types of pattern similarity analyses were conducted: (1) Movie vs. recall within participant: The stimulus-induced pattern (elicited during movie) was compared to the recollection pattern (elicited during spoken recall) for each scene within each participant (Fig. 2A). (2) Movie vs. recall between participants: For each participant, the recollection pattern (elicited during spoken recall) of each scene was compared to the stimulus-induced pattern (elicited during movie) for that scene averaged across the remaining participants (Fig. 2D) (3) Recall vs. recall between-participant: For each participant, the recollection pattern (elicited during spoken recall) of each scene was compared to the recollection patterns for that scene averaged across the remaining participants (Fig. 3A). The preceding analyses each resulted in a single brain map per participant. The average map was submitted to the shuffling-based statistical analysis described above and the FDR-corrected p-value was plotted on the brain for every voxel. The same pattern comparison was performed in the PMC ROI and the results plotted for each individual participant. For the fourth type of analysis, movie vs. movie between-participant, for each participant the stimulus-induced pattern (elicited during movie) of each scene was compared to the stimulus-induced patterns for that scene averaged across the remaining participants. The average R-value across participants was plotted on the brain for every voxel (Fig. S2).

#### Classification of individual scenes

We computed the discriminability of neural patterns for individual scenes during movie and recall (Fig. 4) in the posterior medial cortex (PMC) ROI (same ROI as Fig. 2C, 2F, 3C). Participants were randomly assigned to one of two groups (N=8 and N=9), an average was calculated within each group, and data were extracted for the PMC ROI. Pairwise correlations were calculated between the two group means for all 50 movie scenes. For any given scene (e.g., scene 1, group 1), the classification was labeled “correct” if the correlation with the matching scene in the other group (e.g., scene 1, group 2) was higher than the correlation with any other scene (e.g., scenes 2–50, group 2). Accuracy was then calculated as the proportion of scenes correctly identified out of 50. Classification rank was calculated for each scene as the rank of the matching scene correlation in the other group among all 50 scene correlations. The entire procedure was repeated using every possible combination of two groups sized N=8 and N=9. Statistical significance was assessed using a permutation analysis in which, for each combination of two groups (73 possible), scene labels were randomized before computing the classification accuracy and rank. Accuracy was then averaged across the 73 combinations, for each scene the mean rank across the 73 combinations was calculated, and this procedure was performed 1000 times to generate null distributions for overall accuracy and for rank of each scene. Classification rank p-values were corrected for multiple comparisons over all scenes using FDR.

All above analyses were identical for movie and recall except that data were not extant for all 50 scenes for every participant, due to participants recalling 34.4 scenes on average. Thus, group average patterns for each scene were calculated by averaging over the extant data; for 45 scenes there were data available for at least one participant in each group, considering all 73 possible combinations of participants into groups of N=8 and N=9.

#### Comparison of movie-vs.-recall and recall-vs.-recall maps

In this analysis (Fig. 5) we quantitatively compared the similarity strength of recall-vs.-recall to the similarity strength of movie-vs.-recall. First, we compared recall-vs.-recall (Fig. 3B) correlation values to between-participants movie-vs-recall (Fig. 2E) correlation values using a paired t-test at every point in the brain. It was necessary to use between-participants movie-vs.-recall comparisons because within-participant pattern similarity was expected to be higher than between-participant similarity merely due to lower anatomical variability. To assess where in the brain the difference was statistically significant, we performed a bootstrap analysis wherein the individual participant values for recall-vs.-recall and movie-vs.-recall were randomly swapped between conditions to produce two surrogate groups of 17 members each, i.e., each surrogate group contained one value from each of the 17 original participants, but the values were randomly selected to be from the recall-vs.-recall comparison or from the between-participant movie-vs.-recall comparison. These two surrogate groups were compared using a t-test, and the procedure was repeated 100,000 times to produce a null distribution of t values. The veridical t-value was compared to the null distribution to produce a p-value for every voxel. The p-values were then corrected for multiple comparisons across the entire brain using FDR (q < 0.05) and plotted on the brain (Fig. 4A). This map shows regions where between-participants recall-vs.-recall similarity was significantly greater than between-participants movie-vs.-recall similarity, i.e., where the map in Fig. 3B was stronger than the map in Fig. 2E.

Discriminability of individual scenes was further assessed within the regions shown in Fig. 5B (1568 voxels). For every searchlight cube underlying the voxels shown in Fig. 5B, we asked whether each participant’s individual scene recollection patterns could be classified better using a) the movie data from other participants, or b) the recall data from other participants. Unlike the classification analysis described in Fig. 4, calculations were performed at the individual participant level (e.g., using each participant’s recollection patterns compared to the average patterns across the remaining participants, for either movie or recall). Mean classification rank across scenes was calculated to produce one value per participant for a) the movie data from other participants, and one for b) the recall data from other participants. A t-test between these two sets of values was performed at each voxel underlying the searchlight cubes (Fig. 5C).

#### Memorability of scenes vs. pattern similarity

For each scene, the number of participants who had successfully recalled that scene was counted. We then extracted data from the PMC ROI and calculated the pairwise between-participants correlation during recall (same analysis as in Fig. 3A-C, except pairwise), as well as the pairwise between-participants correlation between movie and recall (same analysis as in Fig. 2D-F, except pairwise), at the scene level. Pairwise comparisons were used because the mean value of pairwise correlations is not affected by the number of participants (and the number of participants was different across data points in this analysis). We calculated Spearman’s rank correlation for the number of participants who successfully recalled each scene vs. the average between-participants pattern similarity during recollection for each scene (Fig. 6A). The same analysis was performed using a default mode network ROI (see Experimental Procedures, “ROI Definition”). In a control analysis, we calculated Spearman’s rank correlation for the number of participants who successfully recalled each scene vs. the average between-participants movie-vs.-recall pairwise pattern similarity for each scene in PMC (Fig. 6B). In another control analysis, we calculated Spearman’s rank correlation for the number of participants who successfully recalled each scene vs. the average between-participants pairwise pattern similarity for each scene during the movie in PMC (Fig. 6C). In another control analysis, we calculated Spearman’s rank correlation for the number of participants who successfully recalled each scene vs. the average between-participants movie-vs.-recall pairwise pattern similarity for each scene in an ROI combining V1-V4 (Fig. 6D, same ROI as Figure S3).

#### Hippocampal inter-participant correlation and subsequent memory

Using an anatomical hippocampus ROI, a timecourse was created for each participant by averaging all voxels within the ROI during movie-viewing. During movie-viewing, inter-participant correlation (ISC) was calculated as the Pearson’s correlation of each individual’s timecourse with the average timecourse of the remaining participants, as described in ^13^. Hippocampus ISC was calculated separately for each scene in the movie for each participant, and these values were binned according to whether each scene was later remembered or forgotten during the recall session, separately for each participant (Fig. 6E). Values for remembered and forgotten scenes were compared using a two-tailed t-test.

Post-hoc analyses were conducted to examine subsequent memory effects in different divisions along the hippocampal axis. The hippocampus ROI was divided into anterior, middle, and posterior (three sections of equal length), and the same analyses were performed to compare ISC during the movie for remembered vs. forgotten scenes.

#### Visualization of the BOLD signal during movie and recall

In order to visualize the signals underlying our pattern similarity analyses, we randomly split the movie-viewing data into two independent groups of equal size (N=8 each) and averaged BOLD values across participants within each group (“Movie Group 1” and “Movie Group 2”). An average was made in the same manner for the recall data from the same groups of eight participants each (“Recall Group 1” and “Recall Group 2”). These group mean images were then averaged across timepoints and within scene, exactly as in the prior analyses, creating one brain image per group per scene. A midline sagittal view of these average brains during one representative scene (scene 36) of the movie is shown in Fig. 7A. For the posterior medial area outlined by a white box in each panel of Fig. 7A, we show the average activity for fourteen different scenes (scenes that were recalled by all sixteen of the randomly selected subjects) for each group in Fig. 7B-E.

#### Reinstatement in individual participants vs. between participants

We performed a second-order similarity analysis: correlation of representational dissimilarity matrices (RDMs)^30,44^ within and between participants. An RDM was created from the pairwise correlations of patterns for individual scenes in the movie (“movie-RDM”) and a separate RDM created from the pairwise correlations of patterns for individual scenes during recall (“recall-RDM”), for each participant.

As each participant recalled a different subset of the 50 scenes, comparisons between movie-RDMs and recall-RDMs were always restricted to the extant scenes in the recall data for that participant. Thus, two different comparisons were made for each pair of participants, e.g., S1 movie-RDM vs. S2 recall-RDM, and S2 movie-RDM vs. S1 recall-RDM. In total this procedure yielded 17 within-participant comparisons and 272 between-participant comparisons. We calculated movie-RDM vs. recall-RDM correlations, within-participant, in a searchlight analysis across the brain volume (Fig. 8A). The same analysis was performed between all pairs of participants (Fig. 8B).

Due to the differing amounts of averaging in the within-participant and between-participant maps (i.e., averaging over 17 vs. 272 individual maps respectively), we did not perform significance testing on these maps separately, but instead tested the difference between the maps in a balanced manner. Statistical significance of the difference between the two was evaluated using a permutation analysis that randomly swapped condition labels for within- and between-participant RDM correlation values, and FDR corrected (q < 0.05, two-tailed). Thus, averaging was performed over exactly 17 participants for each permutation.

#### Reinstatement at a finer temporal scale

To examine movie-vs.-recall pattern similarity at individual timepoints (as opposed to at the scene level), we first extracted data from the PMC ROI. For each scene for a given participant, the pattern corresponding to the first timepoint of recall was compared (using Pearson correlation) to the pattern at each timepoint in the corresponding movie scene. These correlation values were averaged across all scenes and all participants (Fig. S1).

#### Overlap of recollection patterns with visual areas

We examined the overlap between 1) brain areas where recollection patterns were significantly similar across participants (same map as Fig. 3B), and 2) visual areas commonly studied in the literature, by plotting both on the same surface (Fig. S3). Retinotopic visual areas were taken from a probabilistic atlas^54^. Face-selective areas were generated using Neurosynth^66^ with the search term “faces ffa” and thresholded at Z = 7.

*Between-participants pattern similarity in PMC, scene-by-scene*. We calculated between-participants correlation values for individual scenes in the posterior medial cortex (PMC) ROI (Fig. S4). For each scene, each participant’s neural pattern was compared via Pearson correlation to the pattern for the same scene averaged across the remaining participants, then these correlation values were averaged.

---

**Table S1.**
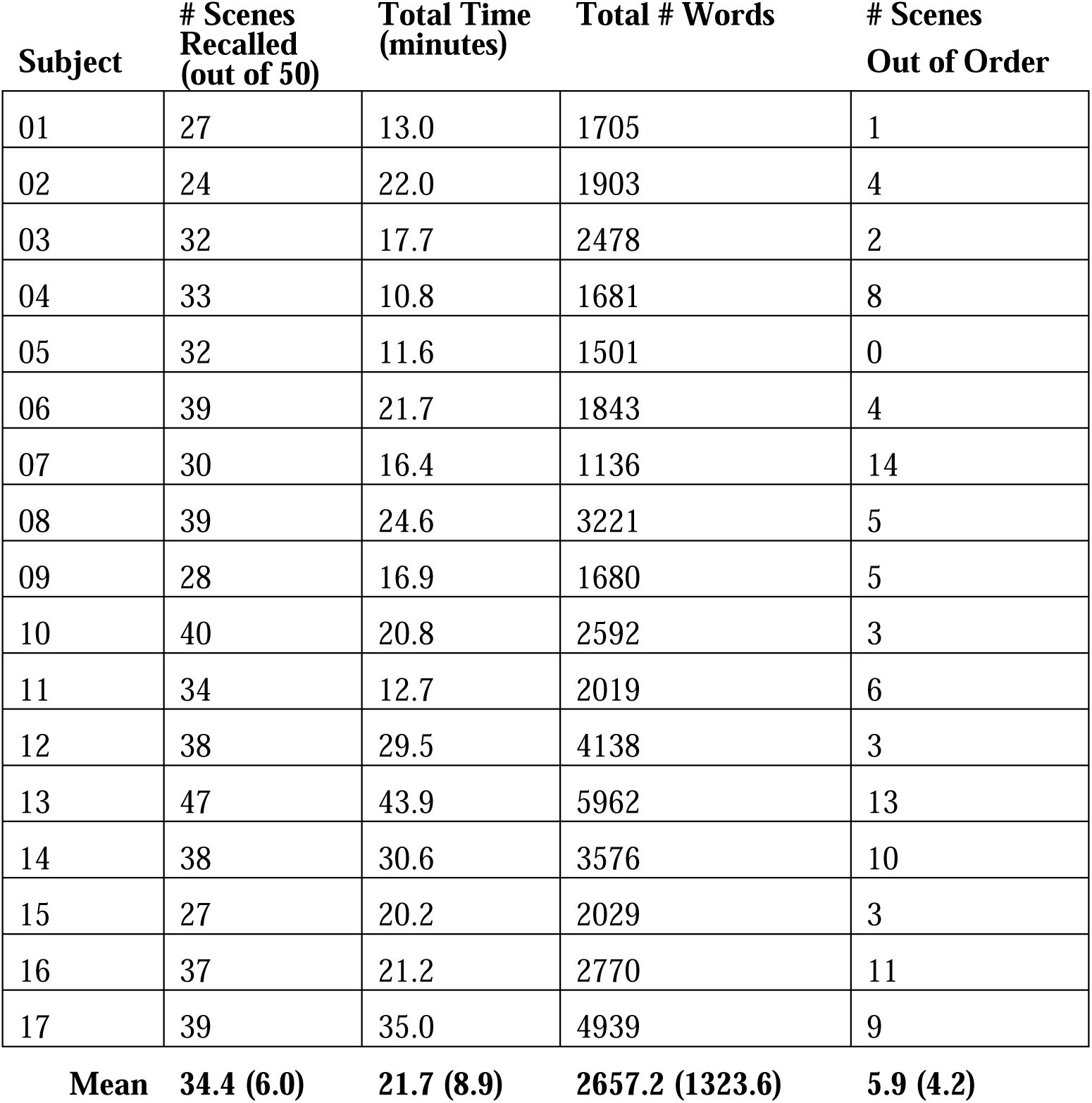
Behavior during spoken recall.

**Table S2.** Examples of four different participants’ descriptions of two movie scenes.

|  |  |
| --- | --- |
| **Scene 13**<br>Duration of original scene:<br>2 min 11 s | **S06:** So then later, at some point in the show, there is a press conference with the Police Chief. The press is asking about the deaths. The chief says they are suicides but feels that they are some how connected. Everyone receives a text that says wrong. The police chief goes on to talk about the deaths. |
|  | **S08:** So we switch to this press conference where the chief of police, or the head of the police station, is answering these questions regarding the fact that these suicides have happened and there are these mass texts that are going on. During this press conference he is talking about the fact that all these suicides are connected in some way because of the similarity of the fact that they have all taken this medicine and they all are in places that they don't normally belong. And these text messages start occurring during the press conference as he's explaining the case that simply say "wrong" to all of the reporters. And he gets a text message that says he knows where to find him and its signed SH, so from Sherlock Holmes. |
|  | **S15:** Yeah so then there's a scene with a detective and he's giving a press conference to some reporters, and its the detective and a woman who he's with. And during the press conference the reporters are asking him questions and as he's answering them, periodically there are texts that pop up and the say "Wrong!" and the reporters are all kind of taken aback. And people giving the press conference instruct the reporters to ignore the texts, but as the press conference goes along, it happens three different times and the third time the head inspector gets a text and its an invitation from SH to have him come seek his help with apparent suicides. During the press conference the reporters are asking if the suicides are actually, or like how they could be being investigated by homicide detectives? The detective doesn't really know much but he's just saying that they think they're all linked because all the people kill themselves the same way and they're all in odd locations when they do it, and there's no suicide notes. |
|  | **S17:** So then we get the press conference, and the guy's saying that he thinks that these suicides are linked, and then a reporter says but how could suicides be linked? that doesn't make any sense. And then the head detect-, the head of the, sergeant or whatever says something like, I don't know but we're investigating it. Then everyone in the room gets a text saying Wrong, and everyone's kind of confused about it, and he said, the woman beside him said If you just got a text please ignore it. And then everyone's asking more questions, and then the sergeant says something that apparently Sherlock Holmes thinks is wrong, because they all get a text saying it's wrong again. |
| **Scene 36**<br>Duration of original scene:<br>1 min 55 s | **S06:** Watson he walks home and he hears a pay phone ring when a third one rings he answers. The voice says look at security cameras across the street and then asks him to get in the car. |
|  | **S08:** So he's walking down this main road and the phone booth next to him rings. And then he continues walking, kind of ignoring that, and another phone rings in a business as he's walking by. And then it stops as someone else comes to answer it. And then he keeps walking, there's a third one that's on his left, and its ringing. So he goes into the phone booth, he answers, and this voice on the phone directs him to see the fact that there are these multiple security cameras that have been kind of tracking where he is. And it tells him to get into this car, to come where he's going. |
|  | **S15:** So Dr. Watson is on his way back and he's passing telephone booths and first one rings, he ignores it, second one rings, he ignores it, third one rings, and he finally answers, there's a voice on the other end that tells him to look at security cameras in the area and each one gets averted. And then a car pull up and he says get in, I don't need to threaten you, you already know. |
|  | **S17:** So then we see, right, kinda like across the street from the building where that lady was found dead, there was this red telephone booth. And it was ringing. Which is weird, and then Watson looks at it and hears it ringing but chooses not to answer it. So then he just ends up walking, we see him walking like on a sidewalk, busy sidewalk, and he's looking in a shop window. And there's a telephone ringing again. And then a store clerk is about to answer it but then doesn't. So then he has like a confused face on. And then he keeps walking and then on the third ringing at yet another phone booth, he is again confused and so okay well I'm just gonna answer it. So he goes in and answers it, and this man, whose voice we haven't heard yet, says, Do you see the camera to your left? And so then he kinda, he says Who is this? and the guys says Do you see this camera. So he looks up, and, the camera, so then we see from the camera's view, we're looking at Sherlock in the telephone booth. And then the camera pans over to the street. And then the person on the phone says, Do you see the camera across the street? And then we see the camera moving across the street. We see the camera on Watson's right. And the guy asks Do you see the camera. And so Watson notices all these cameras. And we get a shot of, in each corner of the screen, a view of the street where Watson is. So then the guy says, ok there's gonna be a car, it's gonna pick you up, do you understand the situation you're in? So get in the car, it's gonna pick you up. So then Watson understands the situation he's in and he gets in the car. |

**Figure S1.**
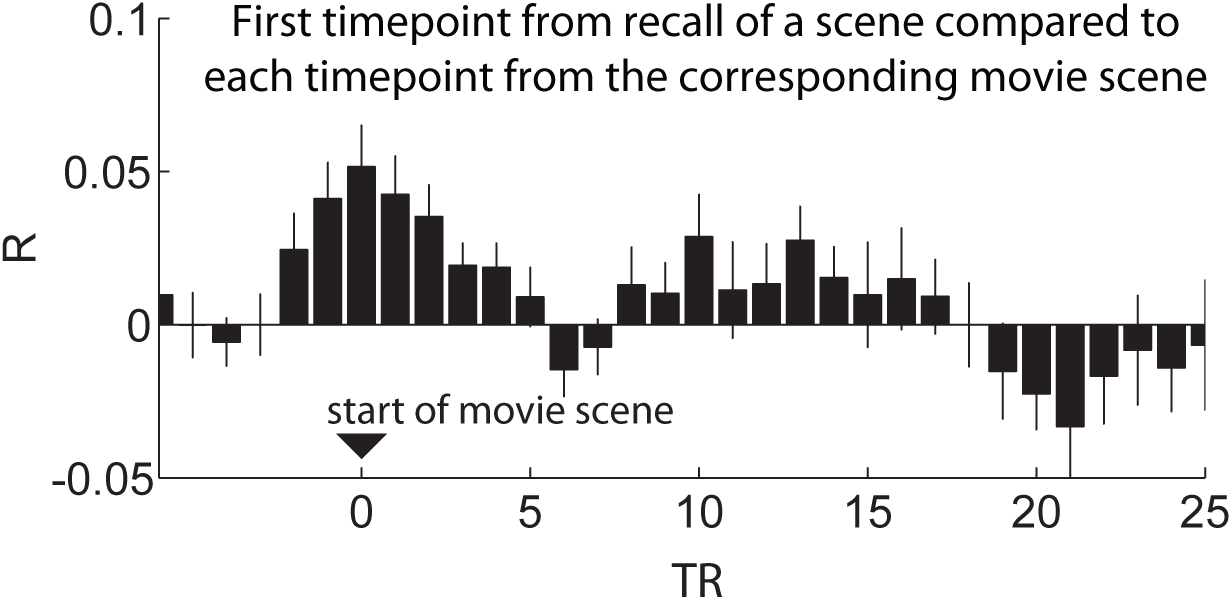
Within-participant pattern reinstatement at a finer temporal scale. While averaging at the scene level was effective for observing neural reinstatement, the behavior of mnemonic recollection that we observed unfolded over time at a finer scale than the scene level. For example, participant 8 used 131 words over 67 seconds to describe scene 13. Here, we further examined reinstatement effects at individual timepoints. For each scene for a given participant, we compared the pattern of activity at each timepoint in the movie scene with that at the first timepoint of recall of that scene in the posterior medial cortex ROI. On average, correlations with the earliest timepoints of encoding scenes were higher than correlations with later timepoints. Error bars represent standard error across subjects.

**Figure S2.**
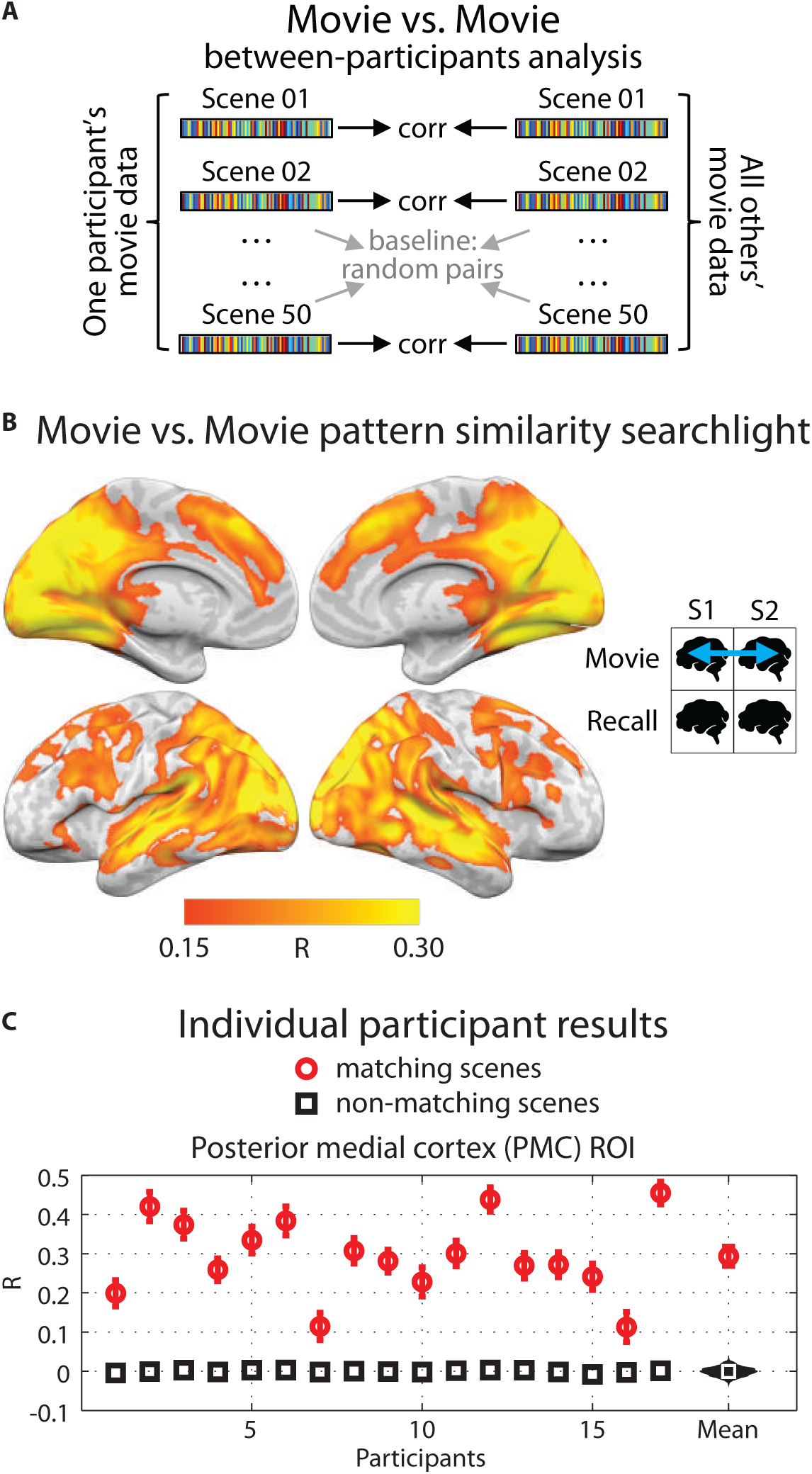
Pattern similarity between participants during the movie. **A)** Schematic for between-participant movie-vs.-movie analysis. BOLD data from the movie were divided into scenes, then averaged across time within-scene, resulting in one vector of voxel values for each movie scene and each recalled scene. Correlations were computed between matching pairs of movie scenes between participants. Statistical significance was determined by shuffling scene labels to generate a null distribution of the participant average. **B)** Searchlight map showing correlation values for across-participant pattern similarity during the movie. Searchlight was a 5×5×5 voxel cube. **C)** Correlation values for all 17 participants in independently-defined PMC (posterior medial cortex). Red circles show average correlation of matching scenes and error bars represent standard error across scenes; black squares show average of the null distribution. At far right, the red circle shows the true participant average and error bars represent standard error across participants; black histogram shows the null distribution of the participant average; white square shows mean of the null distribution.

**Figure S3.**
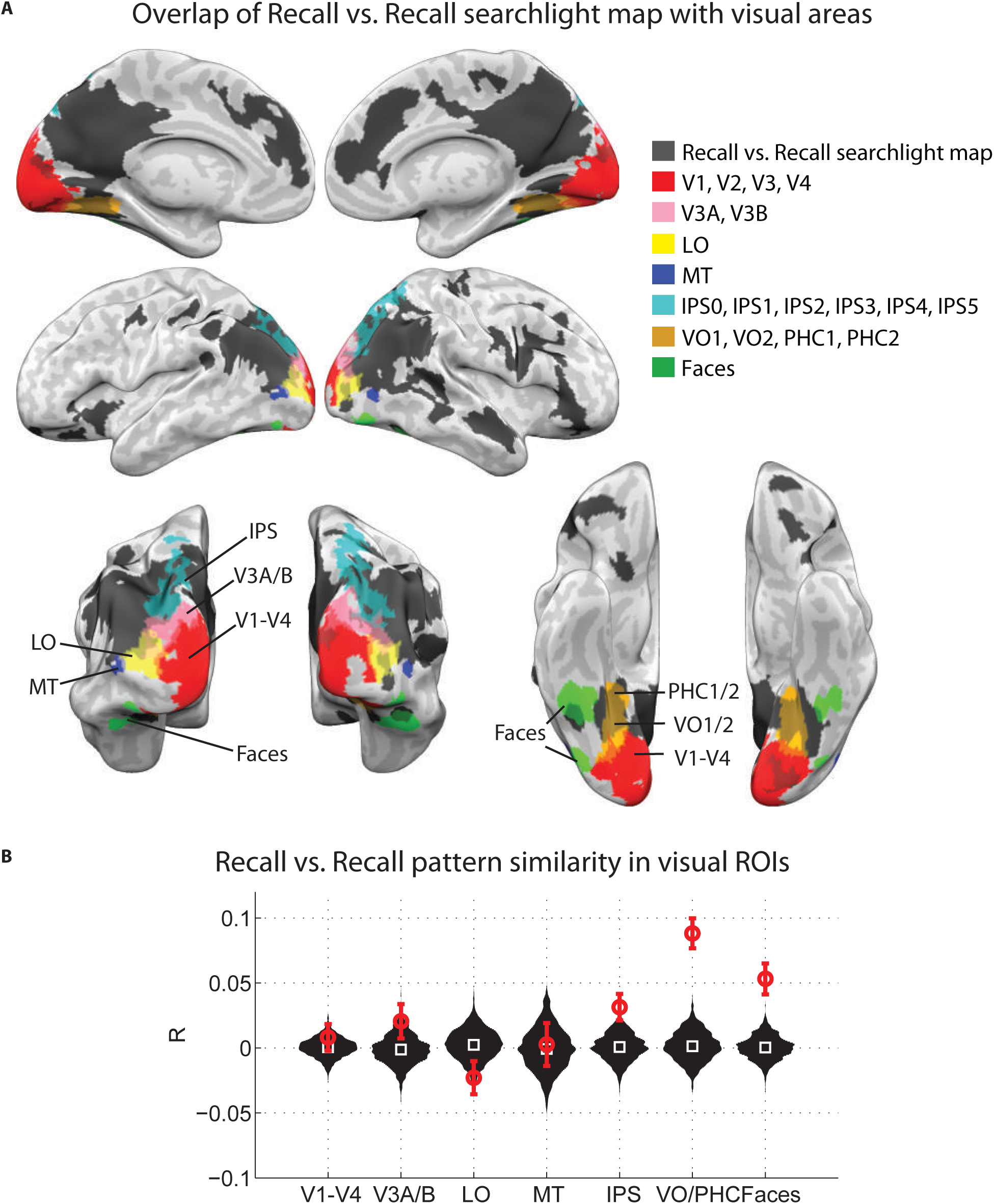
Overlap of recall-vs.-recall map with visual areas. **A)** In gray, brain areas where recollection patterns were significantly similar across participants (Fig. 3B). In other colors, commonly studied visual areas. Retinotopic visual areas were taken from a published probabilistic atlas54. Face-selective areas were generated using Neurosynth66. **B)** For each of the visual area ROIs shown in [A], similarity of scene-level recollection patterns was calculated between participants in the same manner as Figure 3. Statistical significance was determined by shuffling scene labels to generate a null distribution of the participant average. For each region, red circle shows the true participant average and error bars represent standard error across participants; black histogram shows null distribution of the participant average; white square shows mean of the null distribution. In low-order visual regions, recall-vs.-recall pattern similarity was not different from chance; however, significant recall-vs.-recall pattern similarity was observed in higher-order visual regions (VO/PHC and face-selective areas).

**Figure S4.**
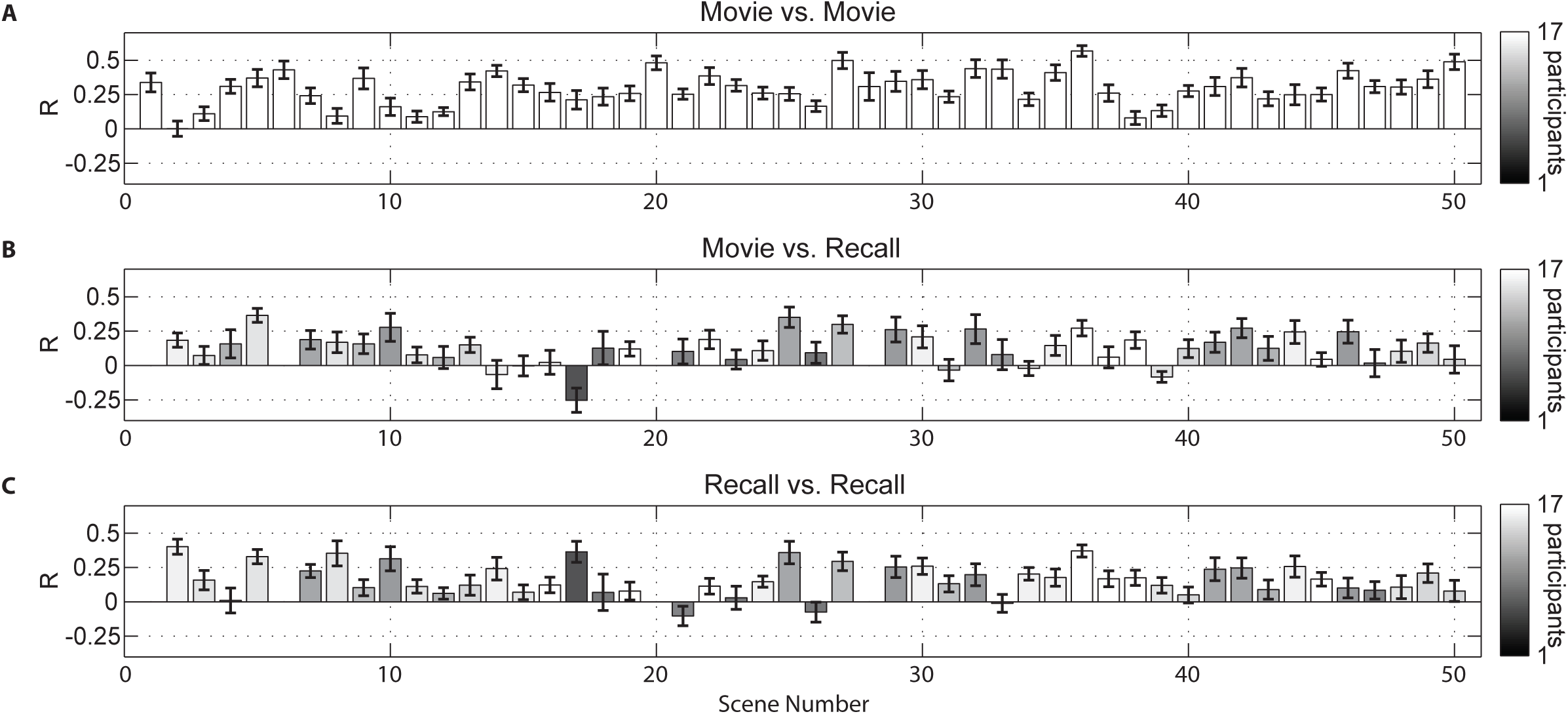
Between-participants pattern similarity in PMC, scene-by-scene. **A)** Between-participants movie vs. movie correlation values for 50 individual scenes in the posterior medial cortex (PMC) ROI (same ROI as Fig. 2C, 2F, 3C). For each scene, each participant's stimulus-induced movie pattern from that scene was compared to the pattern from the corresponding movie scene averaged across the remaining participants. The bars show the average across participants for each scene. Error bars represent the standard error across participants. **B)** Between-participants movie vs. recall correlation values for individual scenes in the PMC ROI (46 scenes were recalled by two or more participants). **C)** Between-participants recall vs. recall correlation values for individual scenes in the PMC ROI.

